# Measuring Metacognition of Direct and Indirect Parameters of Voluntary Movement

**DOI:** 10.1101/2020.05.14.092189

**Authors:** Polina Arbuzova, Caroline Peters, Lukas Röd, Christina Koß, Heiko Maurer, Lisa K. Maurer, Hermann Müller, Julius Verrel, Elisa Filevich

**Affiliations:** Bernstein Center for Computational Neuroscience, Berlin, Germany; Berlin School of Mind and Brain, Humboldt-Universität zu Berlin, Germany; Institute of Psychology, Humboldt Universität zu Berlin, Germany; Neuromotor Behavior Laboratory, Giessen, Germany; Institute of Sport Science, Justus Liebig University, Giessen, Germany; Center for Mind, Brain and Behavior, Gießen and Marburg, Germany; Institute of Systems Motor Science, Universität zu Lübeck, Germany

**Keywords:** voluntary movement, motor metacognition, domain-generality, meta-d’, motor awareness

## Abstract

We can make exquisitely precise movements without the apparent need for conscious monitoring. But can we monitor the low-level movement parameters when prompted? And what are the mechanisms that allow us to monitor our movements? To answer these questions, we designed a semi-virtual ball throwing task. On each trial, participants first threw a virtual ball by moving their arm (with or without visual feedback, or replayed from a previous trial) and then made a two-alternative forced choice on the resulting ball trajectory. They then rated their confidence in their decision. We measured metacognitive efficiency using *meta-d*’/*d*’ and compared it between different informational domains of the first-order task (motor, visuomotor or visual information alone), as well as between two different versions of the task based on different parameters of the movement: proximal (position of the arm) or distal (resulting trajectory of the ball thrown).

We found that participants were able to monitor their performance based on distal motor information as well as when proximal information was available. Their metacognitive efficiency was also equally high in conditions with different sources of information available. The analysis of correlations across participants revealed an unexpected result: while metacognitive efficiency correlated between informational domains (which would indicate domain-generality of metacognition), it did not correlate across the different parameters of movement. We discuss possible sources of this discrepancy and argue that specific first-order task demands may play a crucial role in our metacognitive ability and should be considered when making inferences about domain-generality based on correlations.

## Introduction

Metacognition refers to the ability to monitor and report one’s own mental processes and, due to its ties to theories of consciousness, it has been the object of study of philosophers (Brown et al., 2019; Lau & Passingham, 2006) and cognitive neuroscientists alike (Dehaene et al., 2017).

A common operationalisation of an individual’s metacognitive ability is the relationship between accuracy in a discrimination decision (first-order, or type I task) and subsequent confidence reports (second-order, or type II task). Under this operationalisation, a wide variety of decisions can be subject to metacognitive scrutiny and give rise to a feeling of confidence in their accuracy. Measuring metacognition therefore naturally requires committing to a given domain; because the discrimination judgement must be about, for example, visual, auditory, or tactile perceptual information, or semantic or episodic knowledge. This raises the question of whether a single metacognitive mechanism can monitor all cognitive domains, or multiple, domain-specific mechanisms are needed to monitor each specific cognitive domain. This question is important because it constitutes a first step to understand the cognitive architecture of metacognition. Normally, domain generality is studied by asking the same participants to do two different metacognitive tasks. Domain-generality, then is inferred from the shared variance of measures of metacognitive ability measured in two different domains and across participants. Metacognitive ability has been found to correlate across perceptual tasks (Faivre et al., 2018; Samaha & Postle, 2017; Song et al., 2011), but not between perceptual and memory tasks (Baird et al., 2013, 2014, 2015; Fitzgerald et al., 2017; McCurdy et al., 2013; Morales et al., 2018). Even within memory, different aspects of memory monitoring like prospective and retrospective judgments show dissociations (Chua et al., 2009; Kim & Cabeza, 2009). A recent meta-analysis (Rouault et al., 2018) brought these results together and concluded that metacognitive ability within individuals consistently correlates between different modalities in perceptual domains, but not between perceptual and memory domains. However, Rouault et al. (2019) also noted that this might be due to the differences in the methods and low statistical power. Indeed, a recent study that addressed these problems by having large sample sizes and uniform experimental paradigms found correlations between metacognitive ability across different types of memory, executive function and perception (Mazancieux et al., 2020). In short, it is still not clear to what extent metacognition should be considered domain-general and, more importantly, which aspects of a pair of tasks are predictive of shared variance.

### Motor Metacognition: A Special Case for Metacognition

It has been speculated (Fleming et al., 2014) that the monitoring of internally-generated signals differs from externally-generated ones. Surprisingly, the only non-perceptual domain studied recently is memory (Rahnev et al., 2020) but other domains that rely on internally-generated signals, like motor control, emotions and attention, have not been extensively examined. Motor metacognition represents a very interesting case in the context of this putative internal-external distinction: Voluntary movements are internally generated but, unlike memory, they also elicit rich multi-modal sensory feedback about the executed movement (Haggard, 2005; Wolpert & Ghahramani, 2000). Both the efferent motor command and the afferent sensory feedback might be available to the actor for metacognitive monitoring. And, interestingly, while intuition suggests that there is an introspective, first-person privileged access to internally-generated motor signals, the data so far suggest otherwise. Many motor control and learning processes happen unconsciously or without explicit monitoring (Blakemore et al., 2002); often, only outcomes and the endpoints of the movement are reflected upon (Metcalfe et al., 2013); and a recent study suggested that there is no better metacognitive access to voluntary, active movements as compared to passive movements (Charles et al., 2020).

### Motor Metacognition: The Current State of Research

Early on, Fourneret and Jeannerod (1998) used a reaching task with perturbations to assess awareness of hand movements, and found that participants largely misjudged the effects of distortion of movement direction, as observed in their type I performance. More recently, Augustyn and Rosenbaum (2005) took a different approach: they used a visuomotor reaching task that required participants to optimise their speed-accuracy trade-off. They found that participants were indeed able to find the optimal trade-off given their own performance and suggested that this was the result of successful motor metacognition. However, neither of these studies explicitly collected subjective judgements trial-by-trial, and it is therefore difficult to relate them to recent studies of metacognitive monitoring that adopt the strict discrimination decision followed by confidence judgment design (Fleming & Lau, 2014). Indeed, Bègue (née Sinanaj) et al. (Sinanaj et al., 2015; Bègue et al., 2018) recognised the importance of the differences in operationalisations and studied metacognition of a visuomotor task following this task design. They adapted the seminal task designed by Fourneret and Jeannerod (1998) to create a visuomotor conflict-detection task. Participants reached a target with a joystick and detected deviations from the trajectory introduced by experimenters during the movement. On each trial, they rated confidence in their detection decision. While this task is closer to, and therefore more easily comparable to metacognitive tasks in other domains, it departs from the standard metacognitive task in that the type I task is a *detection*, and not a *discrimination* decision. Lee et al. (2018) showed that the way the type I task is formulated — whether it is a detection or discrimination task — affects measures of metacognition, and suggested that discrimination tasks should be used in order to avoid response biases typical in detection tasks. Moreover, recent evidence suggests that confidence judgments following discrimination and detection decisions rely on partially different neural signatures (Mazor et al., 2020).

In a recent study, Charles et al (2020) aimed at disentangling the contributions of different sources of information in motor metacognitive judgments. They compared metacognition of voluntary finger movements to corresponding passive movements and a visual replay of the movement. Charles et al. (2020) found a positive confidence bias in the active movement condition as compared to the passive movement condition, but no differences in metacognitive efficiency (metacognitive sensitivity for a given level of performance). They argued that while having additional efferent information boosts the overall feeling of confidence, passive movements can be monitored as accurately as active ones.

### Motor Metacognition: Bridging the Gap

However, it remains unknown how different parameters of movement can be metacognitively monitored and to what extent. In the present study, we address this question. We had the following goals: first, in Experiment 1, we developed a paradigm for a metacognitive motor task that is based on a naturalistic movement and we compared it with a visual metacognitive task on the one hand, and a visuomotor metacognitive task, on the other hand. Our pre-registered hypothesis was that there would be no correlation between metacognitive ability in purely motor and in purely visual conditions. We reasoned that this would be the case because the monitoring of voluntary motor movements relies primarily on internally-generated signals, whereas visual monitoring relies on externally generated information alone, and previous literature has suggested that these two general domains might depend on different mechanisms. Second, in Experiment 2, we investigated different participant’s ability to monitor different parameters of movement: a more direct and proximal parameter based on the position of the effector (the forearm) and a more indirect distal parameter of movement based on the effect of the movement (trajectory following a ball throw). We hypothesised that metacognitive access to proximal and distal movement parameters relies on similar mechanisms, and that their relationship to visual metacognition is similar. Additionally, we explored whether metacognitive access to proximal parameters of movement is better than to distal parameters.

Methodologically, for our type I tasks, we opt for a discrimination task across all modalities, to avoid aforementioned effects of a detection task on decision bias (Lee et al., 2018).

## Experiment 1

### Methods

This study was pre-registered (Experiment 1: https://osf.io/kyhu7/), and unless stated otherwise, we followed the pre-registered plan.

#### Participants

Forty participants completed Experiment 1 (29 females, 11 males, mean age: 26.85, range 18 to 36 years; our pre-registered plan was to exclude participants over 35), but the number of the participants included in each of the analyses changed following the pre-registered exclusion criteria. The resulting sample size is specified in each analysis. All participants had normal or corrected-to-normal vision, were right-handed and had no history of psychiatric or neurological disorders (all self-reported). All of them spoke good German or English. Participants were naive to the purpose of the study, and all experimenters were aware of the hypotheses of the study. All participants provided written informed consent before starting the experiment, and were reimbursed for their time and effort with 8€/hour. The procedures were approved by the ethics committee of the Humdoldt-Universität zu Berlin Institute of Psychology and are in line with the Declaration of Helsinki.

#### Apparatus

Participants used a bespoke manipulandum: a metal bar that pivoted around a vertical axis (i.e., in the horizontal plane). At the proximal end of the metal bar (placed just below the elbow, on the vertical axis) a goniometer (Novotechnik RFC4800 Model 600, with 12 bit resolution, corresponding to at least 0.1° precision) measured the angle of the metal bar. On the opposite end, at the tip of the metal bar, there was an electrical switch, which worked similarly to a touch sensor. Analog data were transferred through a Labjack T7 data acquisition device (LabJack Corp., Lakewood, CO) sampling at 1000 Hz. The visual stimuli were displayed on an LCD monitor (2560 x 1440 pixels, 61 cm x 34.5 cm, refresh rate of 60 Hz), placed at approximately 50-60 cm from the participant.

#### Procedure

Experiment 1 consisted of two metacognitive tasks: a classical visual perception task and the novel *Skittles* task. Half of the participants started with the visual task, and the remaining half started with the *Skittles* task.

##### Visual Metacognitive Task (*Dots* Task)

On each trial, two circles filled with dots appeared briefly (200 ms) to the left and right of a fixation cross *(Dots* task; Figure 1A). One of the circles contained exactly 50 dots, whereas the other one contained less or more (between 1 and 100 dots). Participants discriminated (in a 2-alternative-forced-choice task, 2AFC) which of the two circles contained more dots (type I discrimination) by pressing left or right key on the keyboard. After each response, participants used the mouse to report their confidence in their response on a continuous vertical scale from “unsure” to “sure”, and clicked to confirm their type II choice. The starting point for the mouse pointer (displayed as a horizontal bar) on the scale was randomly determined for each trial. First, participants did several trials with feedback to familiarize themselves with the task (until they reported to feel comfortable doing the task). For these initial trials, the cursor on the confidence scale turned green for correct or red for incorrect responses in the type I task. Then, participants performed 40 calibration trials, without confidence ratings or feedback. The difficulty of the type I task (i.e., the difference in the number of dots between the two circles) was adjusted using an online 2-down-1-up adaptive staircase (Leek 2001), aimed at reaching ~71% correct responses in order to find a good starting difficulty level for the main part of the task. Finally, participants performed the main part of the task, with no feedback and the same 2-down-1-up online staircasing procedure. There were 200 trials in this main part.

**Figure 1.**
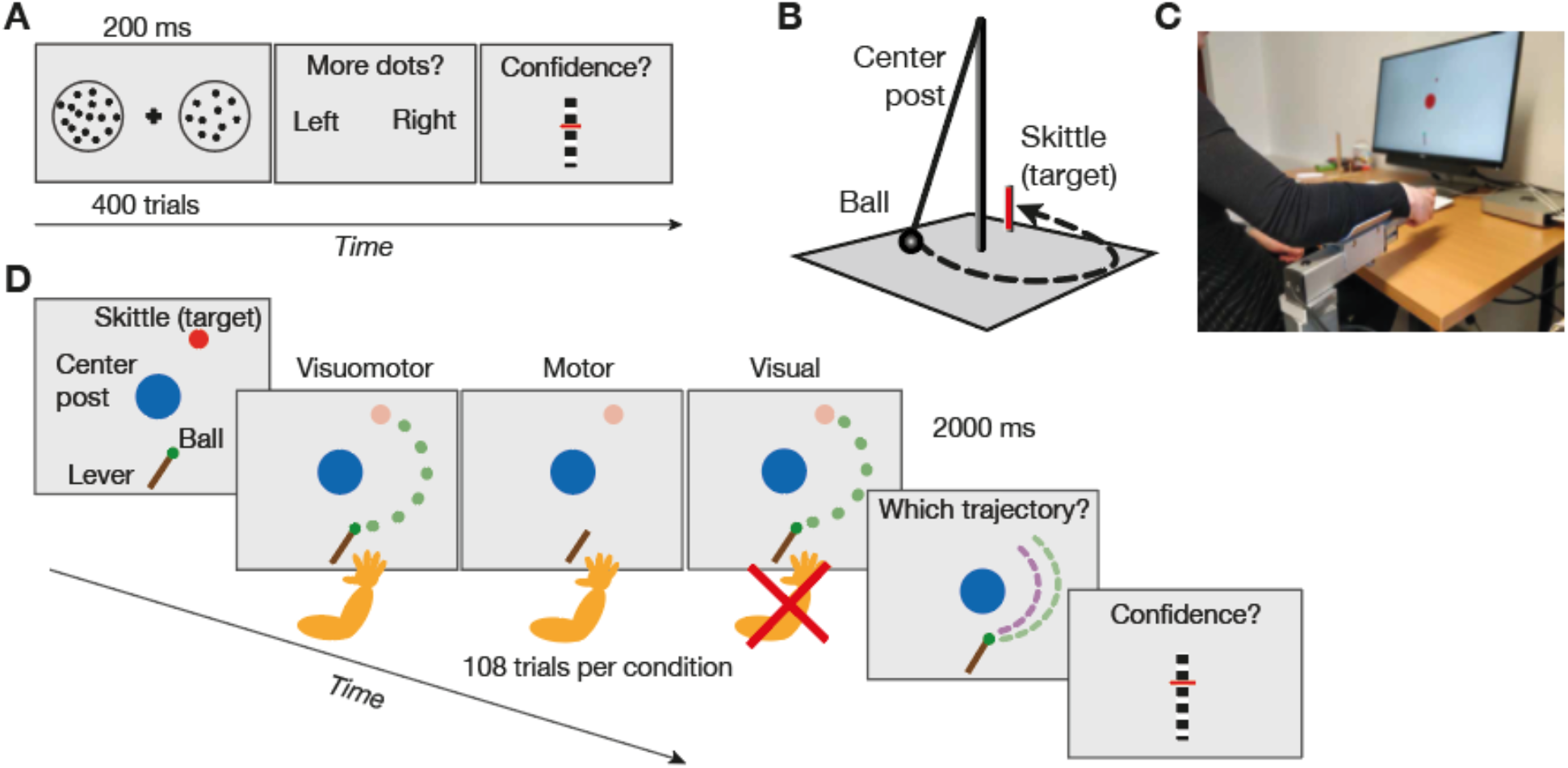
Experimental setup and paradigms. **A.** Paradigm of the visual *Dots* task. On each trial, two circles with different numbers of dots were briefly shown and participants decided which circle contained more dots. Then, they rated their confidence in this decision. **B.** *Skittles* game, the real-life prototype of our motor *Skittles* task. **C.** Setup for the *Skittles* task with the manipulandum and the *Skittles* scene on the screen. **D.** *Skittles* task paradigm. On each trial (in one of the three possible conditions: visuomotor, motor or visual), participants threw a virtual ball and then chose which one of two trajectories displayed on the screen corresponded to their own ball throw. They then rated their confidence about their preceding choice. The ball was shown flying in visuomotor and visual conditions. The dashed lines correspond to the two complete ball trajectories (displayed statically). The target is depicted in lower contrast to illustrate that it was present at the beginning of each trial and disappeared after the ball release.

##### Metacognitive Skittles Task

We built our task on the *Skittles* task (Müller & Sternad, 2004), which is based on a real-world game, where participants swing a ball that hangs from a post to hit a target (a skittle) standing behind it (see Figure 1B). In our computerized version, participants had a bird’s-eye view of the corresponding scene.

Each participant completed three different experimental conditions, namely visuomotor, motor and visual. Participants started each trial of the visuomotor condition with their arms resting on the horizontal metal bar of the manipulandum (see Figure 1C). To “pick up” the virtual ball, participants placed their right index finger on the touch-sensitive electrical switch on the distal end of the metal bar. Participants then accelerated the ball by moving their forearm with the metal bar around its vertical axis and “released” the ball by lifting their index finger from the distal end. This movement effectively mimicked a naturalistic ball throw. Importantly, also, this naturalistic movement can be described with only two parameters: given a set of constants describing the physical properties like the ball mass and radius, the angular velocity at the moment of release and the angle of release fully determine the ball trajectory (for the formal description of the model see Müller & Sternad, 2004 and Sternad et al., 2011). Participants threw the virtual ball in this way, aiming to hit the skittle. We chose the position of the skittles slightly asymmetrically to the right relative to the manipulandum and the central post, to make it easier to hit it with a rightward throw. The flight of the ball was displayed for 600 ms after it had been released. After that, two ball trajectories were displayed on the screen: One of them corresponded to the trajectory of the participant’s own ball throw, whereas the other trajectory differed from the participant’s own in the velocity at the moment of ball release. The exact difference in velocity was determined by an online adaptive staircase, as we describe further. Participants identified in a 2AFC which of the two trajectories corresponded to the one that they had generated. Participants made their choice using the metal bar: One of the trajectories was highlighted (the order of highlighting the correct and incorrect trajectory first was randomized) and participants could change the highlighted trajectory by moving the metal bar (the highlighting changed every 20° of the movement and irrespective of the direction of the movement, in a randomized way). In this way, we avoided the strict mapping between the position of the metal bar and the trajectory chosen to avoid response biases due to low-level motor priming. Participants confirmed their choice by touching the sensor at the distal end of the metal bar. After that, they lifted their hand from the metal bar and reported their confidence in their choice on a continuous scale using the mouse with their right hand, in the same way as in the *Dots* task.

The three conditions of the *Skittles* task varied in the kind of information that was available to participants for the 2AFC (Figure 1D). In the *visuomotor condition,* participants performed the ball throw and saw the ball during its flight. In the *motor condition,* participants threw the ball, but it disappeared upon release. In the *visual condition,* participants passively observed the replay of their own visuomotor trials (400 ms before the ball release to 600 ms after the ball release) without moving their arm. Visuomotor trials from the previous active block were presented in a pseudorandomized order. In all conditions, the target disappeared from the screen at the moment of ball release, to prevent participants from using information about target hit during the 2AFC decision. Participants were instructed to try to hit the target while paying attention to the trajectory of the ball.

Each participant completed three blocks of 36 trials each per condition (108 trials per condition in total). Visuomotor and motor trials were interleaved and pseudorandomized. The colour of the centre post served as a cue for the condition. Visual trials were blocked and occurred after a block of 72 active (visuomotor and motor) trials.

Before starting the main part of the task, participants had 8 trials with feedback to familiarize themselves with the task. They had the chance to repeat these trials until they felt comfortable with the task. After that, a longer calibration part of 50 trials started (visuomotor only), with no confidence ratings. To control the difficulty of the task, we manipulated the difference in release velocity of the alternative trajectory according to a 2-down-1-up adaptive staircase. The starting point of the subsequent experimental trials for all conditions was determined by the last value in the calibration phase. Participants were instructed to use their right arm for both type I and type II responses (executed with the metal bar and mouse, respectively).

In both the *Dots* and the *Skittles* tasks participants had the chance to indicate (by pressing the spacebar) if they had an action slip and gave the wrong response in the 2AFC task (error trial). These trials were not taken into account for the online staircasing procedure and were excluded from further analysis.

#### Analysis

To quantify metacognitive ability, we used the signal detection theory (SDT)-based measure m-ratio *(meta-d’/d’),* as described in Maniscalco and Lau (2012). We used the sum of squared errors (SSE) method to find the *meta-d’* estimates that best fit the data, using MATLAB scripts retrieved from http://www.columbia.edu/~bsm2105/type2sdt/. This method requires discrete confidence ratings. We therefore transformed participants’ confidence ratings on a continuous scale to a discrete 6-point scale by using quantile ranks.

We examined correlations using the Robust Correlation toolbox for MATLAB (Pernet et al., 2013, retrieved from https://sourceforge.net/projects/robustcorrtool/). This procedure prevents outliers from driving correlation values, thereby decreasing the false positive rate. It also provides 95% confidence intervals (CIs) for the estimates based on bootstrapping, which allows to evaluate the precision of the estimate without assumptions about the underlying distribution. We used the skipped correlation function to find the robust data cloud and exclude bivariate outliers based on the box-plot rule.

We used JASP (version 0.11.1) for Bayesian analyses. We followed the categories outlined in Andraszewicz et al. (2015, based on Jeffreys, 1961) to interpret Bayes factors (BFs). BFs 1–3 were interpreted as anecdotal evidence for the alternative hypothesis, 3 – 10 as moderate, 10–30 as strong, 30–100 as very strong, and BFs > 100 as extreme evidence. The same numbers used as denominators in fractions of 1 (1/3, 1/10 etc.) defined the corresponding thresholds for the null hypothesis. The Robust Correlation toolbox does not provide Bayesian statistics. To estimate BFs for the correlation analyses we first cleared the data from outliers detected by the Robust Correlation toolbox and then imported it into JASP. We used the default Cauchy prior for t-tests with a 0.707 scale factor and a default stretched beta distribution with a width of 1 for correlations. All multiple comparison t-tests were Bonferroni-corrected.

#### Exclusion Criteria

As per the pre-registered plan, for each participant, we excluded any individual conditions if the type I accuracy in these tasks was under 60% or over 80%. In the *Skittles* task, we excluded three participants in visuomotor condition, four participants in the motor condition, and two in the visual condition. No participants in the *Dots* task were excluded based on these criteria.

We excluded individual trials if the reaction time in the type I task was under 300 ms or over 8 s. In the *Skittles* task, four participants always responded within this time range. Thirty-six remaining participants, who had too fast or too slow reaction times in the type I task, only had a few trials like this (median: 1.1%, range 0.31%–9.9%). In the *Dots* task, 34 participants had no trials excluded; and others had few trials excluded (median: 0.50%, range 0.50%–2.5%). We also excluded trials if they represented an artificially easy type I task. This was the case when one of the two trajectories in the visual or visuomotor condition hit the pole and the other one did not. The median number of such trials per participant was 18.5% (range 3.7–79.6%) in visuomotor condition and also 18.5% (range 3.7–79.6%) in visual condition.

For each participant, we divided confidence into two bins. We then excluded individual conditions from a participant’s data if they had extremely low or high values for type I or type II hit rates or false alarm rates, (<0.05 or >0.95), as such extreme values do not allow a stable estimation of SDT measures (Barrett et al., 2013; Bor et al., 2017). Although we initially included this exclusion rule in the pre-registration for Experiment 2 only, we applied it to both experiments. In the *Skittles* task, additional 14 participants were excluded in the visuomotor condition, two in the motor condition, six in the visual condition, and one in the *Dots* task. We acknowledge a high number of participants excluded based on this criterion, particularly in visuomotor condition, and address it in the *General Discussion: Response bias* section.

Eight participants did not report any error trials. For other participants, the proportion of error trials was also low (median: 1.39%, range from 0.46% to 5.56%), so no participant was excluded due to the number of error trials (>10%). Because we excluded individual participants and conditions, the number of data points differed for the different correlation analyses. We report the corresponding sample size along with each set of statistics in the *Results* section. The raw data and analysis scripts are available at https://osf.io/kyhu7/.

### Results: Experiment 1

#### Fundamental Measures: d’ and Confidence

To understand the basic structure of the data, we first examined performance and mean confidence levels for the visual *Dots* task and each condition of the *Skittles* task.

##### First Order Performance (d’)

First, although the trial difficulty for each condition was governed by an independent online staircase aimed at fixing performance at approximately 71% correct, we found differences in type I performance between conditions in the *Skittles* task (ANOVA: *F*(3,128) = 19.72, *p* < 0.01, *η*^2^ = 0.31, BF_10_ = 5.60*10^9^, compared to the null model). Participants showed consistently lower *d’* values in the motor condition (M = 0.96, SD = 0.28) compared to the visuomotor (t(23) = −8.76, M = 1.29, SD = 0.22, Bonferroni-corrected p < 0.01, Cohen’s *d* = 1.31, BF_10, U_ = 3.96*10^5^) and visual conditions (t(30) = 8.06, M = 1.27, SD = 0.21; Bonferroni-corrected, p < 0.01, Cohen’s *d* = 1.25, BF_10, uncorrected_ (U) = 1.21*10^5^) (Figure 3A). Participants’ mean type I performance in the visual *Dots* task (M = 1.26, SD = 0.10) was statistically indistinguishable from that in the visuomotor and visual conditions in *Skittles* task, although the variance was smaller. This might be due to the higher number of trials and therefore less noisy estimates of *d’*.

##### Response Bias

Analysis of the type I responses revealed a response bias: participants consistently chose the left (inner) trajectory more often than the right (outer) one in visuomotor and visual conditions but not in motor conditions, although the presentation of the correct trajectory was balanced. The median ratio was 1.57 (IQR 1.10) in the visuomotor condition and 1.44 (IQR 0.69) in the visual condition. In the motor condition, the median ratio was 0.90 (IQR 0.63).

##### Mean Confidence Ratings

There was a difference in mean confidence ratings in the *Skittles* task (ANOVA: *F*(3,125) = 8.93, *p* < 0.01, *η*^2^ = 0.18, BF_10_ = 317.18, compared to the null model) (Figure 2B) and it followed a similar pattern as the one we found for *d’*: on average, confidence in the motor condition was lower (t(19) = −3.49, *p* = 0.015, M = 53.14, SD = 12.58, Cohen’s *d* = 1.04, BF_10, U_ = 9.30) than in the visuomotor condition (M = 65.92, SD = 12.08), and also lower than in the visual condition (t(27) = 2.97, *p* = 0.036, M = 61.03, SD = 15.92, Cohen’s *d* = 0.55), with Bayesian analysis showing anecdotal evidence for the difference (BF_10, U_ = 1.52). A t-test showed no statistically significant differences between mean confidence level in visuomotor and visual conditions (t(21) = −2.23, *p* = 0.22) while the Bayesian analysis suggested anecdotal evidence for the difference (BF_10, U_ = 1.65).

**Figure 2.**
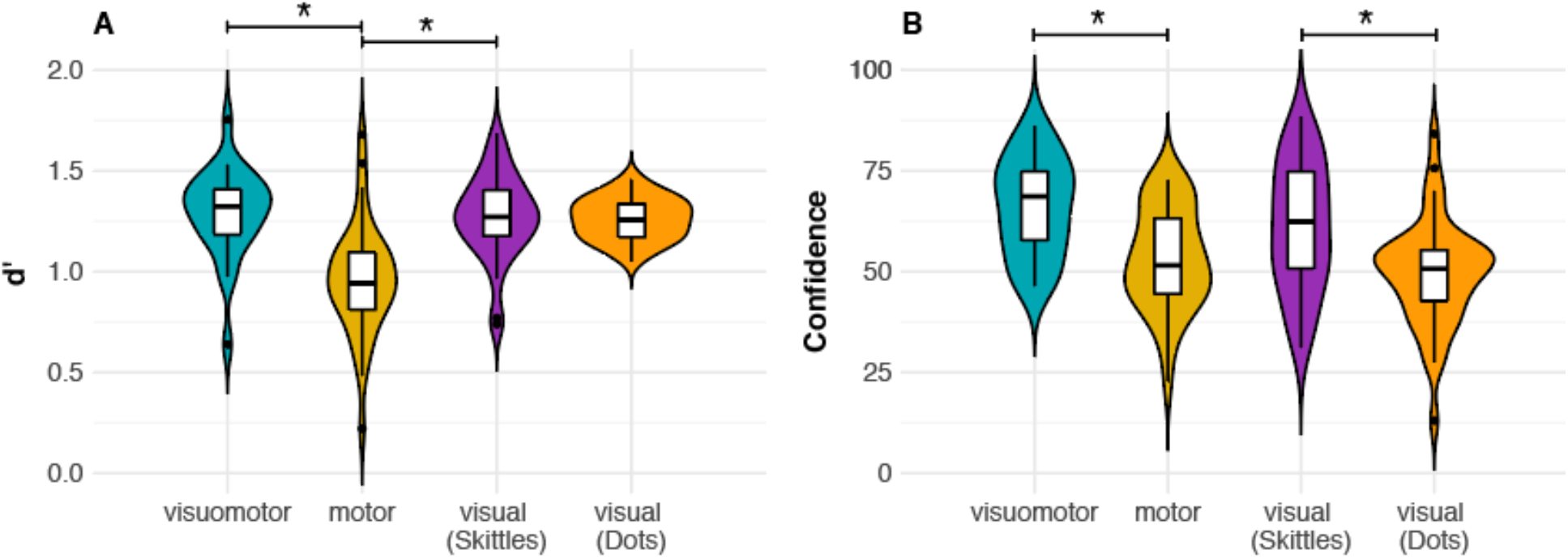
Fundamental measures in Experiment 1. **A.** Mean type I performance and **B.** mean confidence ratings in the three conditions of the *Skittles* and in the visual *Dots* tasks. Asterisks represent significant differences based on *p* < 0.05 (Bonferroni-corrected). The performance in the type I task was lowest in the motor condition of the *Skittles* task. Mean confidence ratings were generally in line with performance differences between conditions, apart from the visual *Dots* task.

Interestingly, while mean *d’* values between the two visual tasks were very similar, confidence was lower in the visual *Dots* task (M = 49.61, SD = 13.32) compared to the visual *Skittles* task (t-test: t(31) = – 4.00, *p* = 0.0022, Cohen’s *d* = 0.78, BF_10, U_ = 13.29). This suggests that despite comparable overall performance, participants perceived the visual *Dots* task as a more difficult task than the visual *Skittles* task.

#### Confirmatory analyses

##### Metacognitive Efficiency: M-ratio

Next, we compared metacognitive efficiency across tasks and conditions. We quantified it as the ratio between *meta-d’* and *d’* (m-ratio). This allowed us to account for differences between conditions in the type I performance. In the *Skittles* task, m-ratio was the highest in the visuomotor condition (M = 0.76, SD = 0.39), followed by the visual condition (M = 0.64, SD = 0.45), and it was the lowest in the motor condition (M = 0.60, S = 0.49). However, the numerical difference in m-ratios was not statistically significant (ANOVA: F(3, 124) = 0.91, *p* = 0.44, *η*^2^ = 0.022, BF_10_ = 0.77): participants’ m-ratios across conditions of the *Skittles* task were statistically indistinguishable from each other. In the same way, metacognitive efficiency in the visual *Dots* task did not differ from that in the visual *Skittles* task (visual Dots task: M = 0.71, SD = 0.27, Cohen’s *d* = 0.19, BF_10_ = 0.31) (Figure 3).

**Figure 3.**
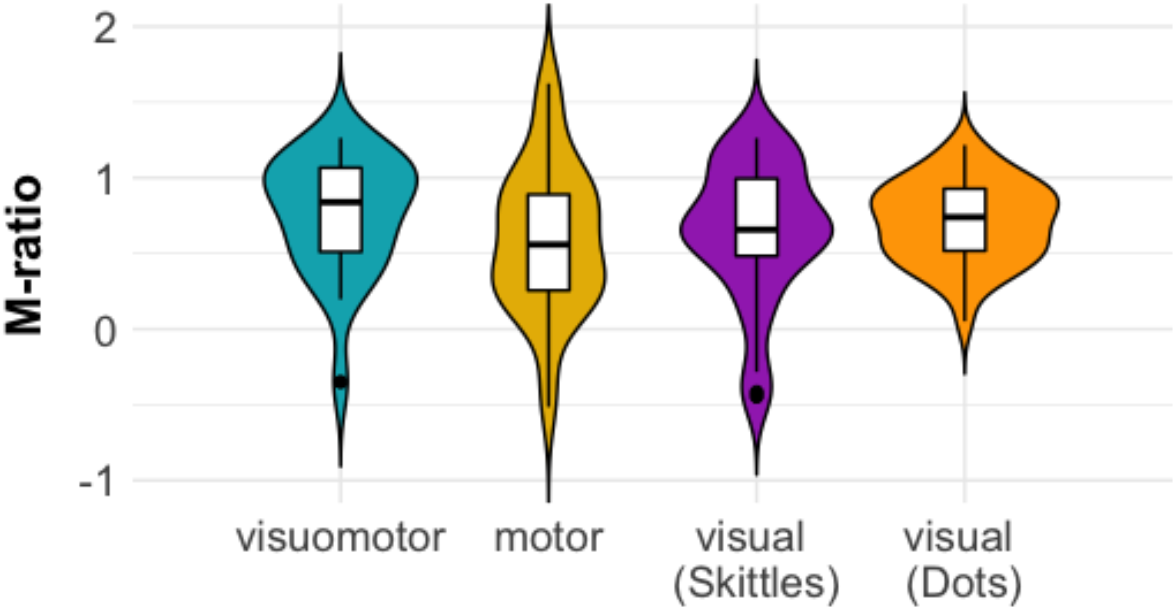
Metacognitive efficiency in Experiment 1. Type II performance measured as *meta-d’/d’* (m-ratio) in the three conditions of the *Skittles* and in the visual *Dots* tasks. There were no systematic differences in metacognitive efficiency between tasks.

##### Correlation Analyses

To directly investigate the relationships in metacognitive efficiency between domains, we ran robust correlation analyses (Pernet et al, 2013) on participants’ m-ratios between different conditions and tasks. First, we compared the two visual conditions (from the *Dots* and *Skittles* tasks, respectively) (Figure 4A). There was moderate evidence for a positive correlation between m-ratios in the visual *Dots* task and those in the visual condition of the *Skittles* task (Pearson’s r = 0.43, CI = [0.09 0.68], n = 31, BF_10_ = 3.45). This shared variance points to an underlying common mechanism that at least partially contributes to (visual) metacognitive processes in both tasks.

**Figure 4.**
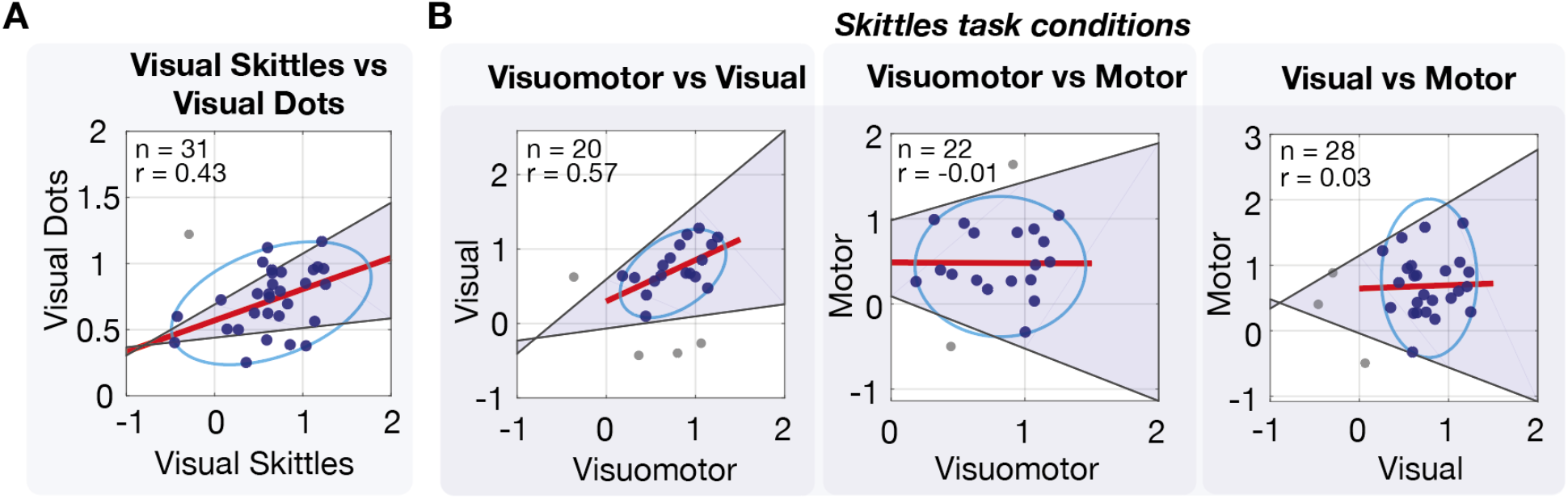
Correlations between conditions and tasks in Experiment 1. Robust correlation analyses between metacognitive efficiency (m-ratios) in the different conditions of the *Skittles* task and the *Dots* task. Confidence intervals (CIs) were obtained via bootstrapping and are shown as shaded areas. Ovals indicate robust data clouds. **A.** Correlation between m-ratios in the visual conditions of the *Skittles* task and the *Dots* task, respectively. Metacognitive efficiency correlates in the two visual tasks. **B.** Correlations in m-ratios between visuomotor, motor and visual conditions in the *Skittles* task. Metacognitive efficiency correlates only between the visuomotor and visual conditions.

We then examined the correlations in m-ratios between the different conditions of the *Skittles* task (Figure 4B, right panel). We found no evidence for a positive correlation between motor and visual conditions (Pearson’s r = 0.03, CI = [−0.37 0.42], n = 28). Bayesian analysis yielded BF_10_ of 0.25, which indicates moderate evidence for the null hypothesis of no correlation. These results suggest that there is no overlap between the mechanisms underlying the monitoring of visual and motor domains. But, because the sample sizes in Experiment 1 were relatively low for correlation analyses, we interpret these results with caution and designed Experiment 2 to examine these correlations in greater detail.

#### Exploratory Analyses: Correlation Analysis with the Visuomotor Condition

We also analysed correlations in m-ratios between the visuomotor condition and its unimodal counterparts, namely the visual and motor conditions (Figure 4B, left and middle panels). This analysis was outlined as ‘exploratory’ in the pre-registration. We found a positive, significant correlation between m-ratios in the visuomotor and visual conditions of the *Skittles* task (Pearson’s r = 0.57, CI = [0.24 0.81], n = 22), BF_10_ = 4.72). This speaks in favour of a common underlying mechanism, presumably driven by a common visual component. In contrast, we found no significant correlation between m-ratios in the visuomotor and motor conditions of the *Skittles* task (Pearson’s r = −0.01, CI = [−0.46 0.47], n = 22), BF_10_ = 0.291). We speculate that the common motor component was too noisy to drive a correlation in this sample, potentially, due to poorer calibration procedure. This, in turn, resulted in a larger drift of the difficulty (see Figure S1). This is corroborated by the significantly larger mean spread of the difficulty (expressed as SD of the velocity) in motor condition as compared to visuomotor condition (t(78) = −4.15, p < 0.001, SD = 0.06).

### Discussion: Experiment 1

In Experiment 1 we compared metacognitive efficiency, measured as m-ratio, on four different tasks and conditions. In three versions of the *Skittles* task, we compared metacognitive monitoring abilities when participants had access to motor-only, visuomotor, or visual-only information. In all tasks and conditions, the vast majority of m-ratios well above zero, which shows that in general, the type II performance is higher than chance and overall, participants knew when they were correct or incorrect. Additionally, and because the *Skittles* task is a novel task, we aimed at validating our results by correlating m-ratios in the visual condition of the *Skittles* task to those from a more standard visual task.

As we expected, we found a significant positive correlation (Pearson’s r = 0.43) in metacognitive efficiency between the two different visual tasks. We note that this was true despite the two tasks being very different in nature. In the *Skittles* task, participants could observe the moving ball’s trajectory for approximately 1 s and were then required to map it onto one of the two trajectories presented on the screen until the decision was made. In contrast, the stimuli in the visual *Dots* task were presented for a relatively short period of time (200 ms) and relied on the representation of the two stimuli in visual short term memory and a numerosity comparison that can happen rapidly and early in the visual system (Park et al 2015). The correlation in m-ratios in these two rather different tasks validate the visual version of the *Skittles* task as a paradigm to investigate visual metacognition.

We then examined our main hypothesis of interest in Experiment 1: We asked whether motor and visual monitoring have any shared variance. Unlike in the comparison of the two visual tasks that we just described (where participants’ confidence was about a visual experience in two formally rather different tasks) we compared participants’ monitoring ability of two different domains across two conditions of the same task. In contrast with the comparison between visual tasks, and despite the strong formal similarities between conditions, we did not find a significant correlation in metacognitive efficiency between the motor and the visual conditions in the Skittles task. Bayesian analysis showed moderate (BF_10_ = 0.25) evidence in favour of the absence of correlation. At face value, this result suggests that metacognition of voluntary movements and visual metacognition rely on different mechanisms. However, we are cautious in committing to this interpretation, as no evidence for correlation might also stem from a relatively low sample size (n = 28), noisy estimate of m-ratios due to low number of trials (the mean number of trials per participant after data pre-processing was: M = 86.8, range 58–101, M = 101.95, range 85–108, and M = 94.5, range 52–108, for the visuomotor, motor, and visual conditions respectively) and drift during staircasing procedure (see S1), which is reflected in SD of the velocity difference: it is significantly larger in motor condition than in visual condition (t(78) = −4.95, p < 0.001, SD = 0.058).

Finally, in exploratory analyses, we found a significant correlation in metacognitive efficiency between the visuomotor and visual conditions but no correlation between the visuomotor and motor conditions of the *Skittles* task. Together, these results suggest that, in the visuomotor condition, participants rely on the visual component more than on the motor one and recall accounts of multisensory integration in situations where the two to-be-integrated sources differ in their precision (Ernst & Banks, 2002; Ernst & Bülthoff, 2004; van Ee et al., 2002).

## Experiment 2

### Methods

#### Participants

Forty-one participants (40 were pre-registered) completed Experiment 2 (29 females, 12 males, mean age: 27.17, range 20–34 years). None of the participants took part in both experiments or any other experiment with a similar task.

#### Apparatus

The same apparatus was used in Experiment 2 as in Experiment 1.

#### Procedure

In the follow-up Experiment 2, participants performed two variants of the *Skittles* task, in two sessions on separate days (not more than 15 days apart, on average 5.17 days apart). Each session lasted approximately 2 hours. In the first variant, as in Experiment 1, participants first discriminated which of two trajectories corresponded to the one they just induced with their movement. We call this task, based on the indirect and distal parameter of the movement, the *Trajectories* task. In the second variant, participants instead discriminated between two possible angles of the arm at the moment of ball release. This decision was based on a more direct and proximal parameter of movement and we call it the *Angles* task (Figure 5).

We introduced a few changes in the *Trajectories* task in Experiment 2 as compared to Experiment 1. First, we increased the number of trials to 200 trials per condition. This yields a more stable estimate of metacognitive efficiency regardless of the fitting method and a reasonably low proportion of false positive results (Fleming, 2017). All trials were split into 5 blocks in each session. Second, participants had more trials to get accustomed to the mechanics of the game and nature of the task: every session started with 8 trials in which participants had to throw the ball and could see the target (without any tasks). This stage was repeated until participants felt comfortable with the game and their throwing. Then they did 8 trials with type I response and feedback. After that, they did 8 trials with type I response, confidence ratings and feedback (as in Experiment 1). Finally, in the initial calibration block with staircasing procedure, both visuomotor and motor trials were used, each with 48 trials (in a pseudorandomized order).

In the *Angles* task, each trial started in the same way and using the same setup as in the *Trajectories* task (Figure 5). However, at the end of the ball flight, participants saw two bars on the screen, representing their arm placed on the metal bar. The angle of one of the bars corresponded to the position of their arm at ball release and the other one was a distractor, rotated clockwise or counterclockwise by a certain angle. The absolute difference between the real angle and the distractor was determined by a 2-down-1-up online staircase (before the start of the main part in the calibration phase and also online -see online supplemental Figures S1 and S2 for individual data from staircasing), and the sign of the difference was pseudorandomized across trials. The position on the screen of the bars representing the correct response and the distractor were pseudorandomized as well. In other words, a distractor bar could either have a larger or a smaller angle than the target bar, and be either on the left or on the right side of the screen. Until the 2AFC, display of visuomotor and visual trials was identical to the *Trajectories* version. In motor trials, we removed the critical visual element: the bar that corresponds to the physical metal bar. The ball was still visible throughout the trial (which was not the case in the *Trajectories* task). As in Experiment 1, the target was present during the preparation of the throw and was removed from the scene after the ball release.

To avoid response biases due to low-level motor priming, participants used a keyboard to report responses in type I and type II questions in both tasks (the “X” and “C” keys corresponded to the left and right response options, respectively). To avoid using more than two input devices, we also used a discrete 6-point scale instead of using a continuous scale (keys “1” to “6” on the keyboard, mapping on to the lowest and highest confidence rating, respectively). Participants were encouraged to use the entire range of the scale. Participants were instructed to pay attention to the angle of their arm at the moment of the ball release, while still aiming to hit the target with their throw (to discourage them from releasing the ball without angular momentum). To keep participants motivated, we displayed the number of target hits at the end of each active block (as they still did not see the target after they released the ball).

#### Analysis

In Experiment 2, we used the same analytical approach and tools as in Experiment 1.

#### Exclusion Criteria

We used the same exclusion criteria in Experiment 2 as in Experiment 1, according to the pre-registration, apart from the reaction times criteria. The pre-registration plan for Experiment 2 stated 200 ms as the lowest threshold, however, for consistency between the two experiments, we used a more conservative threshold of 300 ms stated in the pre-registration of Experiment 1.

First, we excluded participants based on their type I performance (if it fell outside the range 60–80%). This way, in the *Trajectories* task, we excluded two, one and three participants in the visuomotor, motor, and visual conditions, respectively. In the *Angles* task, the performance in all conditions of all participants was within the desirable bounds.

We excluded a low number of trials following the exclusion criterion of reaction times outside the 0.3–8 s range. In the *Angles* task, 27 participants had at least one trial (median: 0.17%, range 0.17%–1.5%) with RTs outside this range, and 14 participants had none. In the *Trajectories* task, 22 participants had at least one excluded trial (median: 0.50%, range 0.17%–4.2%) and 19 participants had none. As per pre-registration, we excluded from the visuomotor and visual conditions in the *Trajectories* task trials in which the real ball trajectory hit the central post and the distractor trajectory did not. One participant had no such trials. The remaining 40 participants had a median 4.5% (range 0.5%–41.5%) of such trials. Visual trials were a replay of visuomotor trials,so the movement parameters (but not the alternative trajectories) were identical. Therefore, the median and range values are the same for both conditions. Based on the criterion of low number (< 5%) of type I or type II hits or false alarms, we additionally excluded five, four, and five participants respectively from the visuomotor, motor, and visual conditions in the *Trajectories* task, and one participant (non-repeating) from each condition in the *Angles* task. In the *Angles* task, 11 participants reported no error trials, and the remaining 30 participants reported a median of 0.25% error trials (range 0.25%–2%). In *Trajectories,* 14 participants reported no error trials and the remaining 27 reported a median of 0.49%, error trials (range 0.25%–3.75%). The raw data and analysis scripts are available at https://osf.io/sy342/.

### Results: Experiment 2

In Experiment 2, each participant completed two variants of the *Skittles* task, on two separate days (in counterbalanced order between participants). On the one hand, as in Experiment 1, each participant completed the three conditions (visuomotor, motor and visual) of the *Skittles* task where they discriminated which of two trajectories shown on the screen corresponded to the one they induced with their movement. Additionally, each participant completed a new version of the task, where they discriminated in a 2AFC manner which of two angles shown on the screen corresponded to the angle of their arm at the moment of ball release. Again, we first checked the overall structure of the data by looking at the type I task measure *d’* and at the mean confidence ratings.

#### Fundamental Measures: d’ and Confidence

##### First Order Performance (d’)

A two-way ANOVA showed a significant interaction effect between tasks and conditions on type I performance (*d*’; F(2,52) = 19.71, *p* < 0.001, *η*^2^ = 0.15, BF_10_ = 1.53*10^13^). Similarly to the results from Experiment 1, pairwise Bonferroni-corrected t-tests revealed that type I performance *(d’)* in the motor condition of the *Skittles Trajectories* task (M = 1.19, SD = 0.21; Figure 6A, left panel) was significantly lower than in the visuomotor (t(26) = 4.84, *p* < 0.01, M = 1.48, SD = 0.30, Cohen’s *d* = 1.12, BF_10_ = 4608.72) and visual conditions (t(26) =3.97, *p* < 0.01, M = 1.46, SD = 0.33, Cohen’s *d* = 0.98, BF_10_ = 1037.00). In contrast, there was no difference between any of the conditions in the *Angles* task (visuomotor: M = 1.17, SD = 0.10, motor: M = 1.20, SD = 0.15, visual: M = 1.16, SD = 0.16; Figure 5 right). Presumably, this was the result of greater similarity between conditions in the *Angles* task due to the fact that participants could see the flight of the ball in all conditions and the attentional demands were better matched than in the *Trajectories* task.

**Figure 5.**
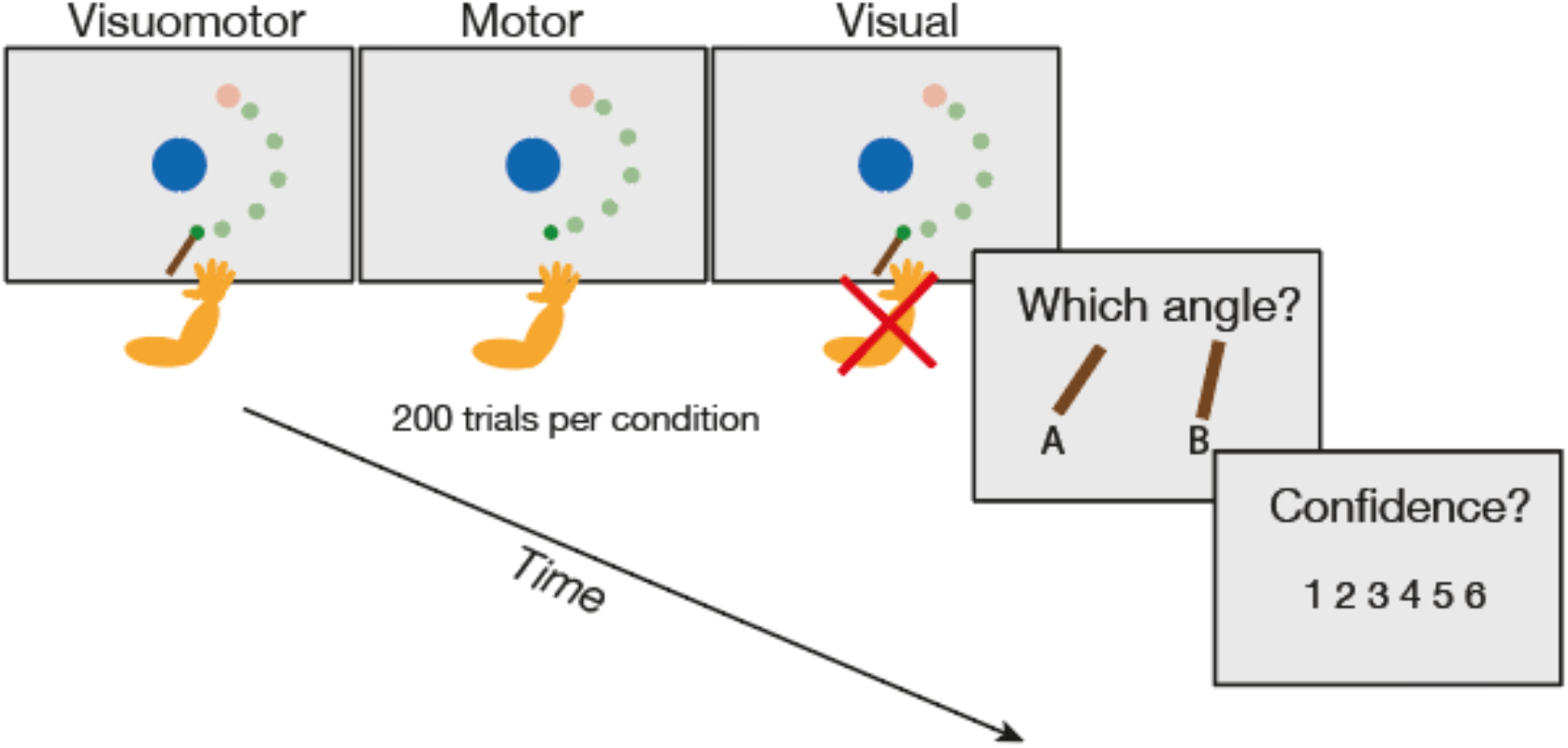
Experimental setup in Experiment 2. *Skittles Angles* task with the 2AFC task about angle of release. We depict a single trial with three possible conditions (visuomotor, motor and visual). Participants threw the virtual ball as in the *Skittles Trajectories* task. They then decided which of two angles displayed on the screen corresponded to their arm position at the moment of ball release. Finally, they rated confidence in their decision. The ball was shown flying in visuomotor and visual conditions. The target is depicted in lower contrast to illustrate that it was present at the beginning of each trial and disappeared after the ball release.

##### Response Bias

In Experiment 2, we observed a similar type I response bias in the *Trajectories* task as in Experiment 1: The median ratio between left and right responses, despite balanced stimulus presentation, was 1.42 (IQR 0.23), and 1.22 (IQR 0.97) for the visuomotor and visual conditions, respectively. As in Experiment 1, responses were more balanced in the motor condition (1.12, IQR = 0.17). However, in the *Angles* task, there was no response bias for choosing the option displayed on the right or left of the screen (ratio between left and right responses, in all conditions: visuomotor: 0.96 (IQR = 0.24), motor 0.96 (IQR = 0.27), visual: 1.11 (IQR 0.61)). That implies that the response bias was not a result of participants simply choosing the option presented on the left or right of the screen. Instead, when we calculated participants’ tendency to choose the larger or smaller angles, regardless of the correct answer (an equivalent of choosing trajectory with larger or smaller velocity), we found a small bias in the motor condition: the ratio between choosing the larger and smaller angle was 1.33 (IQR = 0.24). In the visuomotor and visual conditions, participants did not show a bias (median ratio of responses 1.00 (IQR = 0.14) and 0.90 (IQR = 0.37) respectively). We discuss the response bias further in the corresponding section of the General Discussion.

##### Mean Confidence Ratings

Unlike in Experiment 1, where participants rated confidence in their discrimination response on a continuous scale, in Experiment 2 confidence ratings were expressed on a discrete scale from 1 (“guessing”) to 6 (“very sure”). A two-way ANOVA of the mean confidence ratings in *Skittles Trajectories* revealed a similar pattern of interactions between the tasks and conditions as that of *d’* values across conditions, described above (*F*(2,52) = 26.46, *p* < 0.001, *η*^2^ = 0.072, BF_10_ = 2.75*10^9^) (Figure 6B, left panel). Pairwise t-tests showed that on average, participants were least confident in the motor condition (M = 3.57, SD = 0.62) as compared to the visuomotor condition (M = 4.33, SD = 0.69, t(30) = 9.26, *p* < 0.01, Cohen’s *d* = 1.16, BF_10_ = 2.8*10^9^) and the visual condition (M = 4.05, SD = 0.75, t(28) = 5.99, *p* < 0.01, Cohen’s *d* = 0.70, BF_10_ = 15092.65). Mean confidence ratings were statistically indistinguishable between the visuomotor and visual conditions (t(30) = 5.17, *p* > 0.5, BF_10_ = 0.20). And, again in line with the *d’* results, we found no differences in mean confidence ratings across any of the conditions of the *Angles* task (*p* > 0.5 in all pairwise t-tests; visuomotor: M = 4.07, SD = 0.55; motor: M = 3.97, SD = 0.56; visual: M = 3.90, SD = 0.68) (Figure 6B, right panel).

**Figure 6.**
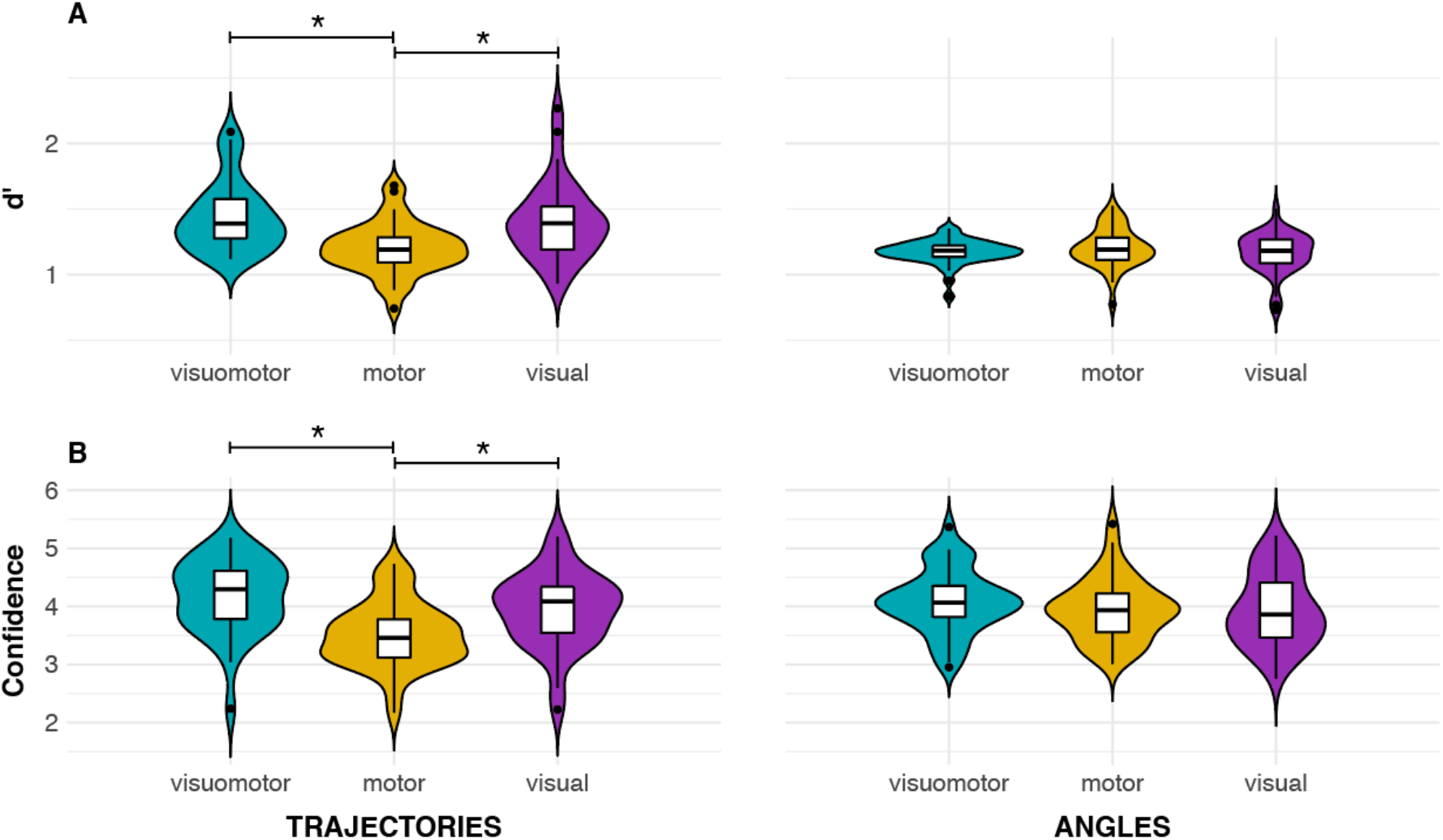
Fundamental measures in Experiment 2. Type I performance (A) and mean confidence ratings (B) in the *Trajectories* and *Angles* version of the *Skittles* task. Similar to Experiment 1 (cf. Figure 2), both type I performance and mean confidence ratings were lowest in the motor condition of the *Trajectories* task. There were no differences in type I performance or in mean confidence ratings between conditions in the *Angles* task.

#### Confirmatory Analyses

##### Metacognitive Efficiency: M-ratio

As in Experiment 1, we quantified metacognitive efficiency using m-ratio. There were no statistically significant differences in m-ratios between any of the conditions or tasks: a two-way ANOVA showed no effect of interactions (F(2,52) = 1.48, *p* = 0.23, *η^2^ =* 0.0083, BF_10_ = 0.016 when compared against the null model) and no main effects of task (BF_10_ = 0.45) or condition (BF_10_ = 0.17), or combination of them (BF_10_ = 0.074) (Figure 7A), although there were differences in *d’* and confidence ratings in *Trajectories.* There were only small numerical differences in m-ratios between conditions (visuomotor: M = 0.79, SD = 0.43; motor: M = 0.64, SD = 0.47; visual: M = 0.87, SD = 0.59). There was no difference in m-ratios between conditions of the *Angles* task (visuomotor: M = 0.58, SD = 0.46; motor = 0.68, SD = 0.45; visual M = 0.61, SD = 0.48) (Figure 7B).

**Figure 7.**
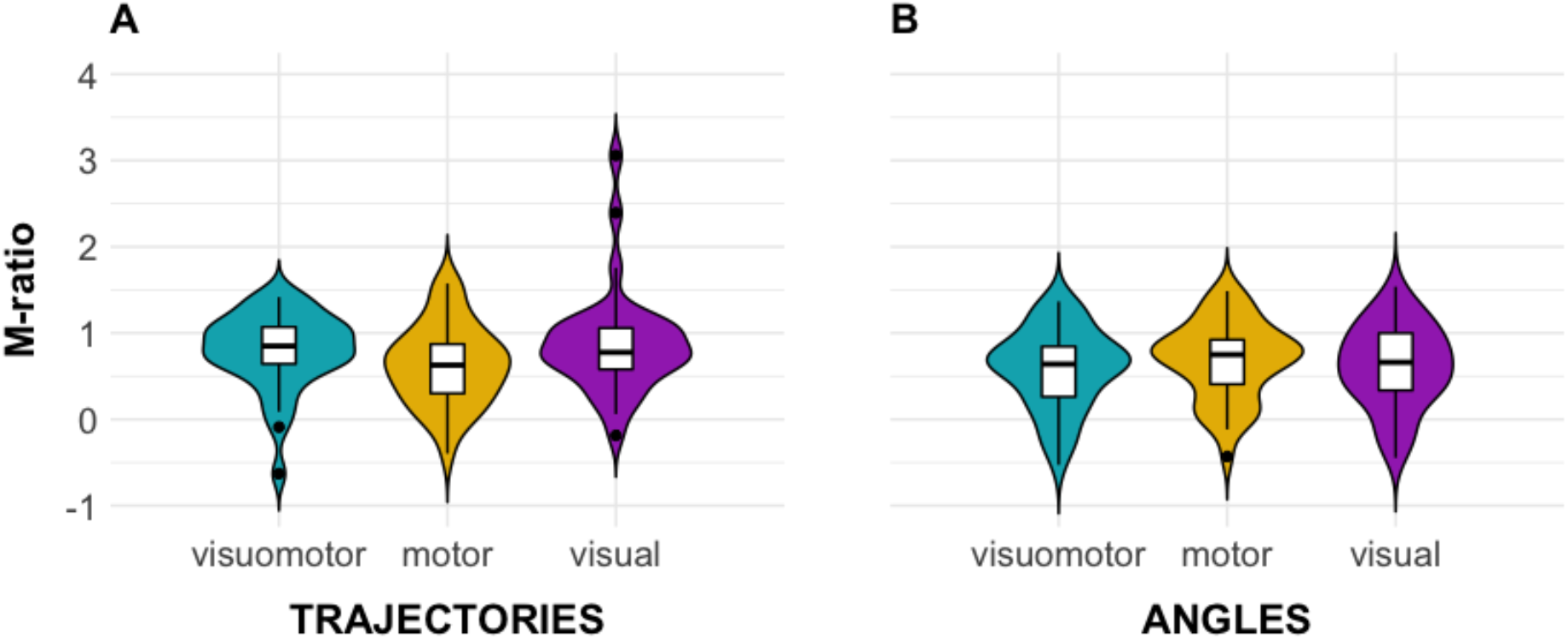
Metacognitive efficiency in Experiment 2. M-ratio across different conditions in the *Trajectories* and *Angles* versions of the *Skittles* tasks. As in Experiment 1 (cf. Figure 3), there were no differences in metacognitive efficiency between conditions.

##### Correlations between Domains

As in Experiment 1, we next sought to investigate the domain-generality of metacognition by examining correlations in measures of metacognitive efficiency between different conditions of each task (Figure 8). In the *Trajectories* task, we found significant, positive correlations in metacognitive efficiency between all conditions. The correlation between the visuomotor and visual conditions, which we observed in Experiment 1 data, was also present in Experiment 2, although it was stronger here (Pearson’s r = 0.73, CI = [0.45 0.86], n = 31 (Figure 8A, cf. Figure 4B, left panel). A Bayesian analysis showed very strong evidence for the positive correlation, too (BF_10_ = 1009.86).

**Figure 8.**
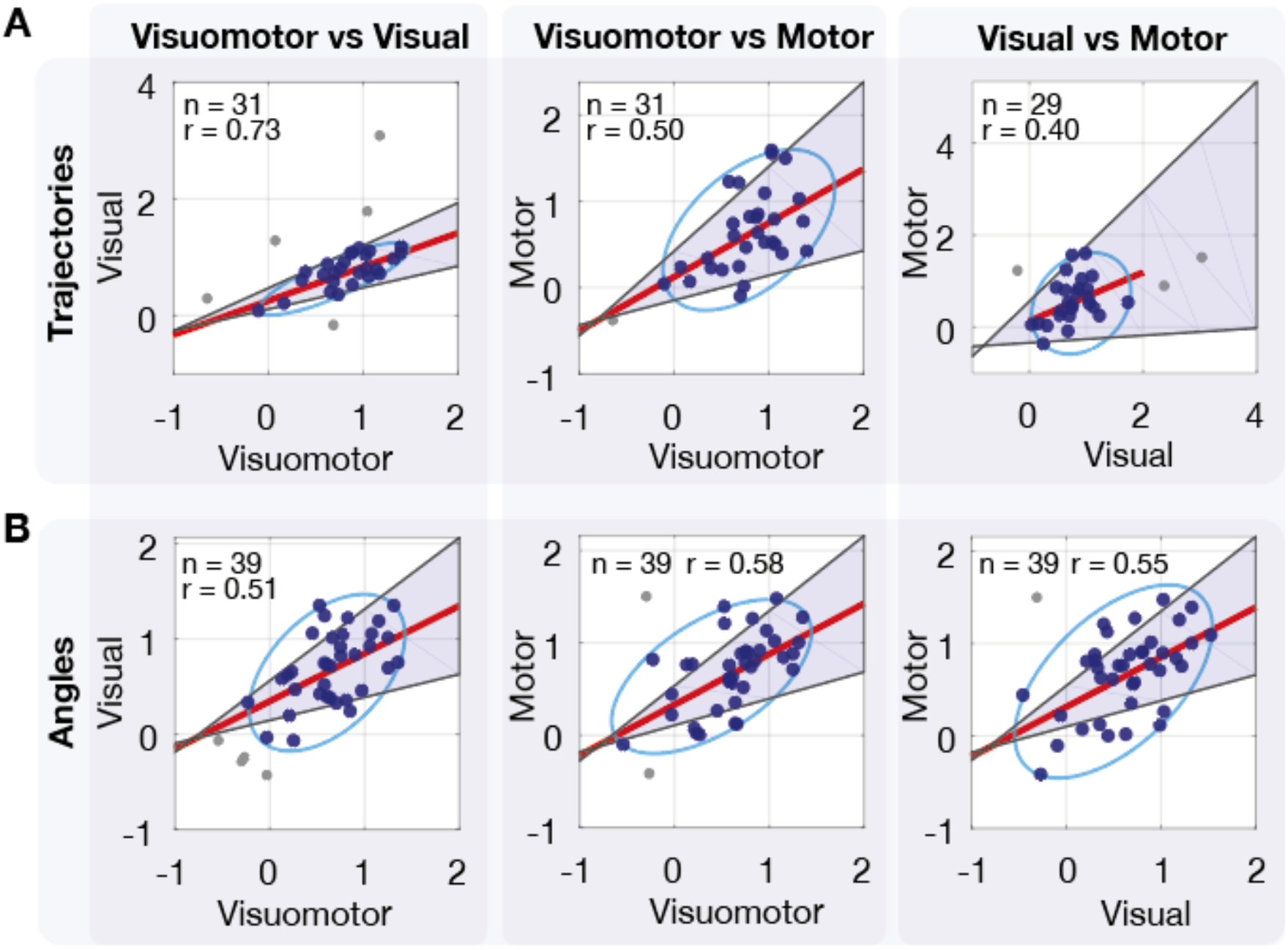
Correlation between different conditions in Experiment 2. Robust correlations between conditions in the *Angles* and *Trajectories* tasks. Confidence intervals (CIs) were obtained via bootstrapping and are shown as shaded areas. Ovals indicate robust data clouds. Unlike in Experiment 1, metacognitive efficiency correlated across all condition pairs, within tasks.

These results from the *Skittles Trajectories* task of Experiment 2 show a somewhat different pattern as those from Experiment 1. First, in Experiment 1 we had found no evidence for a correlation between visuomotor and motor conditions (Fig 4B, right panel). In contrast, in Experiment 2 we did find a correlation between them (Pearsons’s r = 0.50, CI = [0.21 0.69], n = 31, BF_10_ = 9.45) (Figure 8B). More strikingly, whereas in Experiment 1 we found moderate evidence for no correlation between visual and motor conditions (Figure 4B, right panel), we did find a correlation between these conditions in Experiment 2 (Pearson’s r = 0.40, CI = [0.04 0.67], n = 29, BF_10_ = 1.68) (Figure 8A, right panel). We speculate that, the higher number of trials and better staircasing procedure provided better estimates in Experiment 2 for m-ratio in motor condition, as compared to those from Experiment 1. However, the BF_10_ falls within the range of anecdotal evidence and does not allow us to make strong conclusions based on this result.

In the *Angles* task, there were positive correlations in m-ratios between all conditions, too. For visuomotor and visual conditions, it was numerically lower than in the *Trajectories* task (Pearson’s r = 0.51, CI = [0.26 0.72], n = 39, BF_10_ = 24.73) (Figure 8B, left panel). The correlation in m-ratios between visuomotor motor conditions was comparable in the *Angles* task (Pearson’s r = 0.58, CI = [0.34 0.76], n = 39, BF_10_ = 195.39) (Figure 8B, middle panel) to the *Trajectories* task. The correlation in m-ratios between motor and visual conditions was also present, but numerically larger than in the *Trajectories* task and with very strong evidence based on Bayesian analysis (Pearson’s r = 0.55, CI = [0.28 0.73], n = 39, BF_10_ = 98.55) (Figure 8B, right panel), which might stem from less noisy estimates of *d’* in *Angles* and better starcasing in *Angles* task. In fact, difficulty in *Angles* task was significantly less variable than in *Trajectories*, as measured via SD, both in motor (t(80) = −12.42, p < 0.001, SD = 3.37) and in visual conditions (t(80) = −12.66, p < 0.001, SD = 3.62) (see Figure S2).

One of our pre-registered hypotheses stated that the pattern of the correlations between conditions would be similar across the two tasks. In other words, we expected that any correlations between conditions in the *Trajectories* task would be mirrored in the *Angles* task. To test this hypothesis, we treated the three correlations from each task as a correlation matrix and used the Jennrich test: a χ^2^-based statistical test for differences between two matrices (Jennrich, 1970; Larntz & Perlman, 1985). In line with our expectations, we found no evidence for a statistically significant difference between two correlation matrices (χ^2^ = 1.61, *p* = 0.66).

**Correlations between Tasks.** Along with the correlation analysis between different conditions within each task, we also examined the correlations within the same conditions, but *between* tasks. The logic is similar to the one we followed in Experiment 1 to validate the visual condition of *Skittles* task with the visual Dots task: If we assume that these measures capture (at least partially) the same mechanisms, they should correlate across individuals. Strikingly, and against our expectations, we did not find evidence for correlations (Figure 9) in any of the conditions. The confidence intervals of the bootstrapped correlation values between m-ratios included 0 in both the motor condition in the *Angles* and *Trajectories* tasks, and corresponding Bayesian analysis showed moderate evidence for the absence of correlation (Pearson’s r = 0.09, CI = [−0.27 0.40], n = 33, BF_10_ = 0.24), and in the visuomotor ones (Pearson’s r = 0.11, CI = [−0.18 0.45], n = 33, BF_10_ = 0.26). In the visual conditions, although the CI contains zero (Pearson’s r = 0.32, CI = [−0.07 0.64], n = 32, BF_10_ = 0.37) the lower bound of CI for Pearson’s r is very close to zero and the BF lies in the range of anecdotal evidence for the null hypothesis, so this result is more equivocal. These intriguing findings might stem from the differences between the *Trajectories* and *Angles* versions of the task, which we elaborate further in the Discussion section.

**Figure 9.**
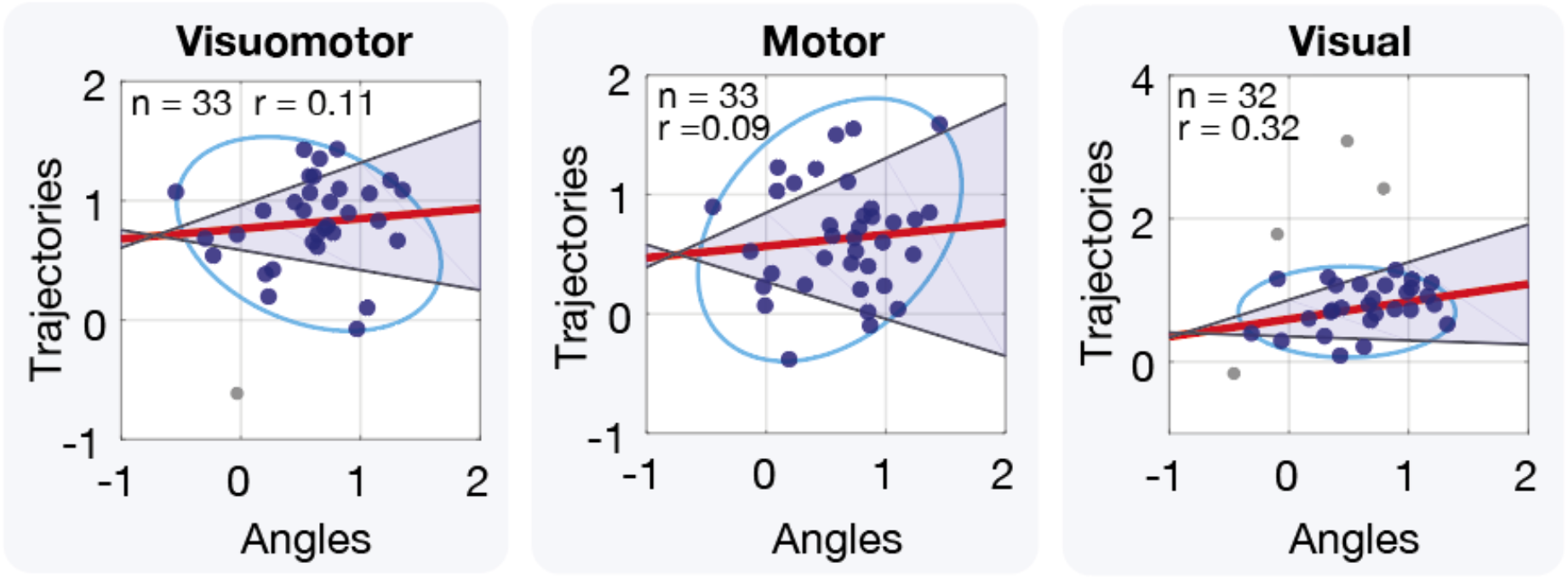
Correlation between different tasks in Experiment 2. Robust correlation analysis for the same conditions across the *Angles* and *Trajectories* tasks. Confidence intervals (CIs) were obtained via bootstrapping and are shown as shaded areas. Ovals mark robust data clouds. In contrast to the correlations between conditions shown in Figure 8, there were no correlations in metacognitive efficiency between the same conditions across two tasks.

#### Exploratory analyses

##### M-ratios in the Motor Condition across Tasks

At pre-registration, we outlined that we would explore relationships between the parameters of movement (the angle of release *vs*. the ball trajectory) and metacognitive ability. We speculated that proximal movement parameters (measured in *Skittles Angles)* would be more readily available to metacognitive monitoring than distal movement parameters (measured in *Skittles Trajectories).* This would lead to higher m-ratios for the motor condition in *Angles* than in *Trajectories.* Contrary to our expectations, when compared across tasks m-ratios in the motor conditions did not differ statistically (t(26) = −0.25, *p* = 0.81); and, in fact, the evidence supports the notion that the two conditions do not differ (BF_10_= 0.18).

##### Motor Behaviour During Visual Trials

Finally, in Experiment 2 we recorded movement of the manipulandum during the visual trials. This allowed us to examine whether participants moved their arms in visual conditions. We confirmed that there was indeed none or very little movement during visual trials (apart from one participant, who moved in 12% of trials in *Trajectories* task and four participants in *Angles* task who moved in 14%, 14.5%, 23.5% and 28% of the trials; their behavioural data were nonetheless included, in order to remain close to the preregistered analysis plan). On average, 2.91% percent of trials had significant movement (>3°) during *Skittles Trajectories* task and 4.38% during *Skittles Angles* task. This suggests that most participants in most rials indeed made their decision based on visual information alone and did not rely on additional information derived from movement execution.

### Discussion: Experiment 2

Apart from the physical constraints of the external world (gravity, air resistance, etc.), the trajectory of a ball after a throw is defined by the movement of the effector at the moment of ball release. In our task, the movement of the effector can be described with two parameters: the angle of the arm and its velocity. Prominent theories of motor control are based on the notion that we control movements in terms of their perceptual consequences (ideomotor theories, Elsner et al., 2002; Hommel, 2013), action goals (Wohlschläger et al., 2003), and facilitated integration of feedback based on distal cues as compared to proximal feedback (as in theory of internal and external focus, Wulf, 2013; Wulf et al., 2002). On the other hand, judging one’s own movement indirectly, based on its effect in the external world (the flight of the ball) relies on a series of mental transformations and understanding of the contingencies between the executed movement and the corresponding visual cues. Therefore, in Experiment 2, we asked whether the monitoring of indirect parameters differed from the monitoring of a more direct and proximal parameter of movement. We asked participants to monitor the angle of their arm at the moment of ball release.

The *Skittles Trajectories* task of Experiment 2 essentially constituted a conceptual replication of Experiment 1, but Experiment 2 differed from Experiment 1 in a series of important methodological aspects, including the confidence scale (continuous in Experiment 1, discrete with 6 levels in Experiment 2), better training and staircasing procedures, and feedback on the number of successful target hits. Similarly to Experiment 1, we did not find significant differences in metacognitive efficiency between different conditions (Figure 7). We also observed the same pattern of results in the confidence ratings in *Skittles Trajectories,* even though we used a discrete confidence scale instead of a continuous one (Figure 6B). As in Experiment 1, in general, participants had good insight of their performance, as the mean m-ratios were well above zero. Despite differences in the mean confidence ratings, participants were not worse at discriminating correct from incorrect 2AFC judgements in the motor condition than in the visuomotor or visual conditions. Thus, our results suggest that metacognitive access to motor information is as precise as that to other sources of information. This goes against early results that suggested that motor monitoring is poor (Fourneret & Jeannerod, 1998).

Most importantly, we found correlations in Experiment 2 (but not in Experiment 1) between visual and motor metacognitive efficiency in *Skittles Trajectories.* The data from the two experiments leads to conflicting conclusions, so the evidence for a positive correlation is equivocal. We speculate that this discrepancy could be due to noisier m-ratio estimates in the motor condition in Experiment 1, due to lower trial count (200 trials per condition in Experiment 2 vs. 108 in Experiment 1) and a better staircasing procedure in Experiment 2. Thus, we argue that this pattern of shared variance revealed in Experiment 2 suggests a common underlying mechanism, which points to some degree of domain-generality across motor and visual metacognition. In both Experiments 1 and 2, we found a positive correlation between the visuomotor and visual conditions. This underlines the salience of the shared visual component in visuomotor and visual conditions in *Trajectories* task. In contrast to Experiment 1, we did find a significant correlation in *Skittles Trajectories* in metacognitive efficiency between visuomotor and motor conditions in Experiment 2. In visuomotor and motor conditions, additionally to higher number of trials, we also had a higher sample size in Experiment 2 than in Experiment 1, thus, a better estimate of the correlation. Taken together, this suggests that the meta-motor component is noisier and has less impact in a multisensory scenario when combined with visual information (like in visuomotor condition).

In the *Angles* task, similar to the *Trajectories* task in Experiment 2, we found positive correlations in metacognitive efficiency across all combinations of the conditions, including between visual and motor conditions, with a BF indicating very strong evidence for correlation (Figure 8B). We therefore take the result from both tasks in Experiment 2 as evidence for some shared metacognitive mechanism supporting visual perception and voluntary movements.

#### Metacognition of Direct and Indirect Parameters of Movement

We expected higher metacognitive efficiency of a more direct, proximal parameter of movement (angle at the moment of ball release) as compared to the metacognition of a more indirect, distal parameter of movement (ball trajectory). Contrary to our expectations, we found no differences in metacognitive efficiency in the motor conditions between the two tasks. In our task, the ball trajectory is defined by the combination of the two release parameters. Since the type I decision in the *Trajectories* task was based on the visually displayed trajectories, participants had to estimate the trajectory by first estimating their velocity and then combining it with the information about the angle (which did not differ between the two trajectories and was displayed on the screen). Despite this additional computational step, participants were as good in reflecting upon their movement in the *Trajectories* version as in the *Angles* version (Figure 6).

One of our pre-registered hypotheses was about the equality of the patterns of correlations between the *Angles* and *Trajectories* tasks in Experiment 2. While there are suggestive numerical differences in corresponding correlations between the two tasks, we found no statistically robust evidence for their inequality. However, it might be due to a low power of this test for small correlation matrices (Larntz & Perlman, 1985). A Bayesian test would be a more suitable alternative here, if it were available.

The most intriguing result is the absence of correlations between corresponding conditions in different tasks. By the same token as in the validation of visual *Skittles* task with visual *Dots* task in Experiment 1, we expected correlations between the same modalities in different tasks (e.g., motor *Trajectories* and motor *Angles).* However, this was not the case: we found no evidence for positive correlations in any of the three conditions (and, in two cases, we instead found evidence for the null hypothesis of no correlation). A trivial explanation to this dissociation is that the two tasks were completed on different days. We argue that this explanation is not likely in the *Limitations* section of the Discussion. Instead, we attribute these dissociations to both the movement parameter that needs to be monitored and deep differences in the task structure. One potential factor could be the difference in the temporal properties between the *Angles* and *Trajectories* tasks. In all conditions of the *Angles* task but only in the motor condition of the *Trajectories* task the critical information needed for the type I task lasted an instant: It was the precise angle of the forearm at the moment of ball release. In the visuomotor and visual conditions of the *Trajectories* task, instead, the evidence that informed the type I decision was available for a more extended period of time (1 s) even if participants had missed the instant of ball release. This difference in temporal features might have interacted with the effects of attention, too. In the more timing-sensitive *Angles* task, effective allocation of attention in time would be more pronounced, whereas in *Trajectories,* momentary attention slips would have a larger effect in the motor condition only. However, if this were a decisive factor, we would expect a correlation in metacognitive efficiency between motor conditions of *Angles* and *Trajectories,* both of which, according to our reasoning, depended on instantaneous information. The fact that we did not find such correlation shows that temporal properties of motor information are not enough to explain the common variance patterns that we observed within the tasks. Instead, it could be a true effect of the parameter being monitored.

## General Discussion

In this study, we asked to what extent shared cognitive mechanisms underlie metacognitive monitoring across two modalities — vision and voluntary movement. This question is important in the context of understanding relationships between the multiplicity of domains of metacognitive monitoring, for several reasons. The first reason is methodological: If we want to make inferences about metacognitive monitoring in general, is it enough to operationalise it with any convenient task? Or would these inferences not generalise to other domains? The second reason is that it may have important societal implications: The ability to correctly monitor our own mental states may be beneficial for efficient information seeking (Boldt et al., 2019; Desender et al., 2018, 2019), education (Zohar & Barzilai, 2013), and in general, for the complex cumulative culture which is characteristic to humans (Dunstone & Caldwell, 2018). But this ability is not static and it may be malleable to training (Baird et al., 2014; Carpenter et al., 2019; Schwiedrzik et al., 2009). Understanding the shared variance between metacognitive domains would help us design training and intervention programs to optimally target any given cognitive function. Previous work measuring the extent of the domain-generality of metacognition has typically assumed that it is the modality of the monitored information that determines which mechanism comes into play. Thus, according to this view, the distinguishing characteristic of a visual metacognitive task is that confidence ratings follow discrimination judgments on visual stimuli regardless of, for example, the speed of evidence accumulation, attention allocation, contextual information, or possible heuristics. Together, our results challenge this view and emphasise the importance of the specific task demands.

### Relationships between Informational Domains

In three datasets (Experiment 1, Experiment 2: *Trajectories,* and Experiment 2: *Angles),* we compared the monitoring of purely motor and purely visual information. We found somewhat inconsistent results, with Experiment 1 showing moderate evidence for no correlation and the two *Skittles* tasks *(Trajectories* and *Angles)* in Experiment 2 showing evidence for a correlation. Because we substantially improved the experimental design in Experiment 2 (increased power, more trials, more comparable staircase procedures across conditions), we consider that the latter results are more robust (but remain cautious in our interpretations). The evidence available in the literature was largely consistent between similar and comparable perceptual tasks, but discrepancies were revealed between perceptual and memory tasks (Rouault et al, 2018). It has been speculated (Fleming et al. 2014) that a distinction may exist between the monitoring of externally generated (i.e., perceptual) and internally-generated information (e.g., memory). By going beyond the classical perceptual and memory tasks, our study provides evidence that the domain-general aspects of metacognitive monitoring might be farther-reaching than previously thought.

### Dissociations between Tasks in Metacognitive Monitoring

Because in our Experiment 2 each participant completed two different variations of the *Skittles* task (*Trajectories* and *Angles*), we had the opportunity to examine correlations in metacognitive efficiency within the same informational domain, but between two different tasks. Strikingly, for all three conditions, we found either evidence for the null hypothesis of no correlation, or no evidence for the alternative. It is, of course, possible that the analysis method of choice is not optimal. Structural equation modelling (SEM), factor analysis or principal component analysis (PCA) might be more powerful tools to reveal common latent variables with differential loadings on observed ones. Future studies of motor metacognition with larger sample sizes may benefit from using SEM to study domain-generality. Alternatively, there might be some intrinsic differences between the tasks that drive the dissociations between tasks and it is our definitions of domains that we have to revise. Instead of only defining them by modality, we should perhaps focus on other aspects of the task, such as the particular temporal properties, attentional demands, or cognitive load, and interactions between them. In line with this argument, Samaha and Postle (2017) showed that the correlation between metacognitive ability in a visual task and a visual short term memory (VSTM) task emerged when the same visual feature was used in both tasks, and not when different features were used. Further, it is increasingly recognised that the type of the task for the type I question plays an important role in metacognitive performance. Lee et al (2019) and Mazor et al (2020) showed dissociations both behaviourally and in the neural mechanisms involved in the computation of confidence following discrimination vs. detection tasks. This might be the reason why the literature on domain specificity yields a somewhat mixed picture (Rouault et al., 2018). In the motor metacognition literature in particular, different studies have alternatively used detection type I tasks (Sinanaj et al., 2015; Bègue et al., 2018) or discrimination tasks (Charles et al., 2020), as in our study. Future studies should carefully consider their type I task design.

We propose that the monitoring of attention (and, consequently, the effective allocation of attention both in time and space) may have been differentially affected by condition in our *Trajectories* and *Angles* tasks. There is little understanding of how exactly attention influences metacognition, although there are indications that it can do so in intricate ways. For example, awareness of visual stimuli is attuned when visibility of stimuli varied in a top-down fashion by varying cognitive load and attention, as compared to more bottom-up visibility changes of contrast or using binocular rivalry (Kanai et al., 2010). Temporal attention modulates confidence, too, as shown in the attentional blink paradigm (Recht et al., 2019). Without a clear understanding of how attention is monitored and controlled, our conclusions remain speculative. Future work may address this interesting direction.

Finally, the rate of evidence accumulation may have driven differences in metacognitive efficiency both between tasks and between individuals. Our task yielded point estimates of performance, insensitive to the temporal dimension of the decision-making process, but emphasising speed over accuracy in the type I task can affect confidence ratings (Pleskac & Busemeyer, 2010; Vickers & Packer, 1982). Differences in the dynamics of post-decisional information accumulation may also affect measures of metacognitive ability (Moreira et al., 2018). In our case, individuals with fast evidence accumulation processes would have shown high metacognitive efficiency for both *Angles* and *Trajectories* tasks, whereas a slow evidence accumulator might have shown higher metacognitive efficiency for the *Trajectories* task (in the visuomotor and visual conditions) as compared to the *Angles* task.

### Absolute Differences between Informational Domains

We assessed the quality of metacognitive monitoring based on motor information alone by comparing metacognitive efficiency in the motor conditions of the *Trajectories* and *Angles* tasks to their visuomotor counterparts. In both cases, we found no differences in metacognitive efficiency between the two conditions. Instead, both of them appeared equally available for introspective insight. Moreover, levels of metacognitive efficiency were statistically indistinguishable from those from the corresponding visual conditions. These results speak against the view that humans have poor access to the parameters of their own movement and predominantly monitor the terminal state of the movement or its effect in the world (Blakemore et al., 2002, Metcalfe et al., 2013). One critical aspect of our *Skittles* task might explain these differences: The focus of the metacognitive task was not on motor performance itself, but on the monitoring of performance. That is, neither the type I nor type II question asked participants directly whether they thought that they had hit the target (although participants were encouraged to try to hit the target during the instructions). Hence, participants may have allocated attentional resources differently in the *Skittles* task than in other motor tasks studied earlier, where motor performance was indeed emphasised. In cases where motor performance is central to the task it might be more beneficial to monitor reaching a goal, or hitting a target, as opposed to monitoring the low-level parameters of movement (Wulf 2013; Wulf et al., 2002). We argue that this distinction is an interesting feature of our results. While very fine and precise motor control can undoubtedly occur in the absence of awareness and attention, we were able to probe the limits of which information about our own movements is *in principle* accessible to metacognitive monitoring.

One important caveat to these analyses of differences between conditions is that m-ratio has a theoretical maximum of 1 (although it often exceeds 1 in practice). Because the mean m-ratios we obtained were relatively close to 1, our null results may be due to a ceiling effect that limited our ability to detect potential differences in metacognitive efficiency between tasks and conditions.

### Response Bias

In the visuomotor condition of the *Skittles* task in Experiment 1,35% of participants of the initial number of participants were excluded based on the criterion that was applied to ensure the stability of SDT-derived measures. One potential reason for such a high exclusion rate could be the response bias that was quite prominent in visuomotor condition (see *Response bias* in *Results: Experiment 1).* In fact, in ten out of 14 participants the ratio of left and right responses was higher than 2. When response bias is high, it is more likely that type I or type II false alarm or hit rates for one of the stimulus are very low or very high. Notably, in Bor et al. (2017), the exclusion rate based on the very similar criterion was comparable: it was 27 from 90 participants (30%) overall, and ranging from 19% to 43% in different subgroups.

In the *Trajectories* task, the response bias was higher in visuomotor and visual conditions, as compared to the motor condition. This suggests that it was driven by the visual components of the task and potentially due to expectations about the behaviour of physical objects (“intuitive physics”, Kubricht et al., 2017).

We also observed a slight bias to choose a larger angle in *Angles* task of Experiment 2. We speculate that this response bias might reflect participants’ beliefs about the point of ball release: during the training phase, some participants reported that they felt that they released the ball at a larger angle than they did, confusing the angle of the ball release with the maximum angle of their arm movement (i.e., the angle at which their elbow stopped extending), which in the rightward movement is always larger than the angle of the ball release. Alternatively, this bias could also be related to the intuitive physics explanation (Kubricht et al., 2017): If participants have biased expectations about the trajectory of the ball (for example, they believe it is more straight than it is), then this can make them believe that the ball was released at a later time point.

### Limitations

Here, we aimed to compare participants’ performance in two tasks and three conditions that differed both in the movement parameter monitored (distal, in *Trajectories,* and proximal, in *Angles)* and the kind of information available (visual, motor, or both). To make the stimuli naturalistic and keep participants motivated throughout the experiment, we made some compromises that resulted in a loss of systematicity in our comparisons. For example, in the *Angles* but not in the *Trajectories* task, participants saw the ball during its flight in the motor condition. Conversely, participants saw the bar on the screen representing the metal bar in the motor condition of the *Trajectories*, but not the *Angles* task. While taxing for participants, a fully factorial design, with the factors corresponding to information about the ball (present or not present), information about the metal bar (present or not present) and the type I question type (about the angle at the ball release moment or about the trajectories) may have allowed us to better identify the reasons for the observed positive correlations (or lack thereof).

Participants completed the different conditions of the same task within the same calibration session, but the two different tasks (*Angles* and *Trajectories*) in two different testing sessions, on average approximately 5 days apart. While it is plausible that short-term fluctuations in metacognitive efficiency led us to find no correlations between tasks, we argue that this is unlikely, as metacognitive ability appears to be a relatively stable trait. Inter-individual differences in computational mechanisms behind the generation of confidence signal remain the same over weeks (Navajas et al., 2017) and metacognitive ability has clear structural brain correlates (Fleming et al., 2010; Fleming & Dolan, 2012; McCurdy et al., 2013; Sinanaj et al., 2015), which are unlikely to change organically, in the absence of training, within days.

### Future Directions

Our results suggest that people can in fact accurately monitor their voluntary movements in the absence of corresponding visual cues. Of course, movement monitoring may in turn involve the monitoring of efferent motor commands, afferent sensory feedback, or a combination of both. Tasks and experimental settings different from the *Skittles* task may help to disentangle the relative contribution of each of these two sources of information to movement monitoring. Charles et al. (2020) aimed at doing exactly this and compared mean confidence and metacognitive efficiency between conditions of active and passive movement, that differed in whether the motor command was present or not. In order to be able to fully study the dissociation, it would be beneficial to have an inverse case, too, where motor commands are present, but proprioceptive feedback is absent. Patients with selective peripheral deafferentiation or volunteers under local anaesthesia may provide such cases. Alternatively, noise can be introduced to proprioceptive feedback, for example, by applying vibration to the relevant muscles and tendons (Fuentes et al., 2012; Goodwin et al., 1972). Studying the effects of the privileged access to motor commands can provide insights for understanding the link between metacognition of voluntary movements and the sense of agency.

While this study did not aim at investigating multisensory metacognition, it is not yet well understood how confidence ratings are computed in multisensory cases. Faivre et al. (2018) compared two models in their ability to describe confidence responses in an audiovisual scenario: an integrative model (two sources of perceptual evidence are first combined to form a confidence response based on the joint evidence) and a comparative model (confidence is computed separately for each source and then combined into one single summary response, for example, by taking the minimum confidence of the two confidence signals). The results from Faivre et al. (2018) were not conclusive and future work may follow this important question that addresses the computational mechanisms underlying domain-general or domain-specific confidence judgments. If the integrative model (as described by Faivre et al., 2018) is valid, it remains an open question if multisensory integration happens in a precision-weighted way, as in type I tasks (Ernst & Banks,2002), or follows different computational principles. Our setup did not allow us to answer this question, as it requires a systematic modulation of signal to noise ratio in one of the sensory channels. However, future research in this direction would be needed to understand multisensory metacognition better.

### Conclusion

We measured human participants’ ability to metacognitively monitor their voluntary movements in a naturalistic task. We found that participants had above-chance metacognitive access to their movements based on either the proximal or on the distal parameters. Further, correlations between informational domains within tasks speak, at face value, in favour of the domain general nature of metacognition. However, the discrepancy we found between two different versions of the task within the same modality underlines the importance of task features when measuring metacognition of a complex process such as voluntary movement. Further work is necessary in order to disentangle contributions of different task features and processes that depend on them (such as attention) to metacognition.

## Context

A rich literature has examined different aspects of motor awareness, including the emergence of the awareness intentions, performance monitoring and perception of learning. Studies on motor awareness most often rely on subjective reports that are known to be subject to biases. In parallel, the field of research on metacognition has grown and developed increasingly sophisticated methods that rid us from precisely this kind of biases. But these two research traditions have remained strongly separated. Here we aimed at bringing them together.

To do that, we had started studying motor metacognition with the *Skittles* task as we describe it in Experiment 1, but we wondered whether the task that relies on participants making a transformation to link the arm movement to the ball trajectory would yield the same results as a different task that did not require this transformation. We therefore modified the original task design and created the *Angles* task to measure a property of movement that was closer to the movement itself, and less so to its effects in the world.

