## Supplemental Material for "Measuring Metacognition of Direct and Indirect Parameters of Voluntary Movement"

**Staircasing**

Here we provide plots from the staircasing procedures. These plots are important for addressing a potential issue with the task difficulty control and unequal variance between task conditions. Data from the calibration period are plotted as negative trial counts and values from the online staircasing procedure are plotted as positive trial counts. If there were more than one calibration block, we plot the last one.

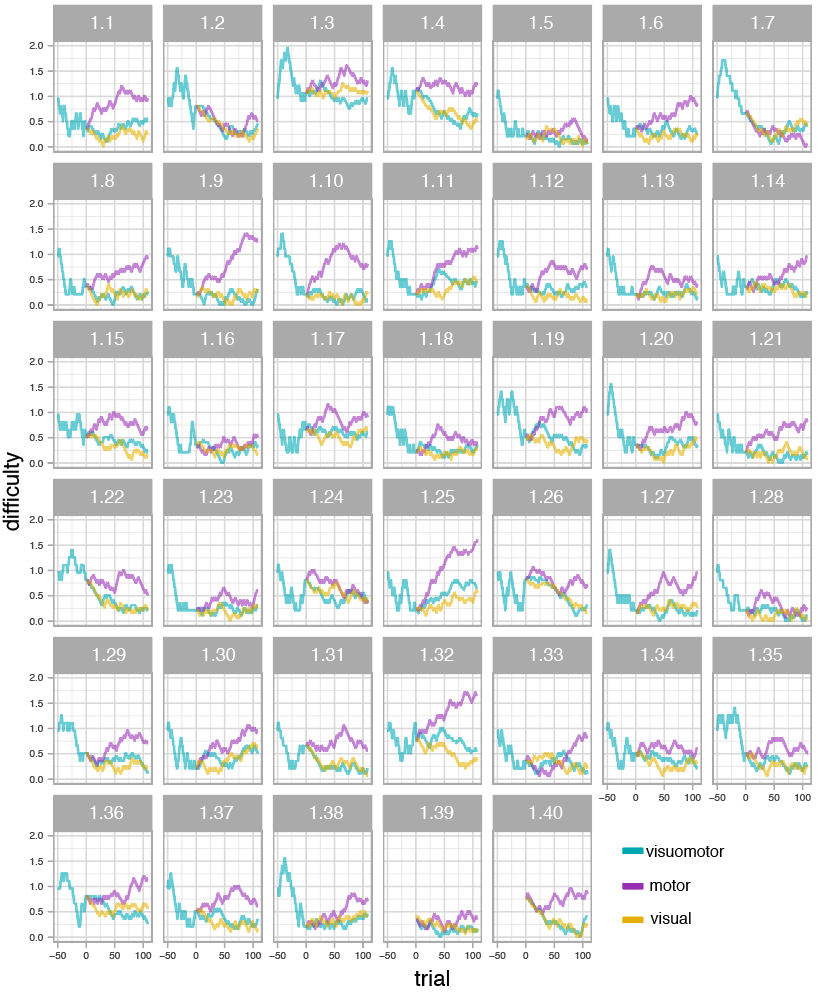

**Figure S1. *Staircasing procedure in Experiment 1, Skittles task.*** Difficulty is expressed as velocity (m/s). Only the visuomotor condition was used in the calibration period. Staircase data for two participants (020SM and 110SR) was not saved.

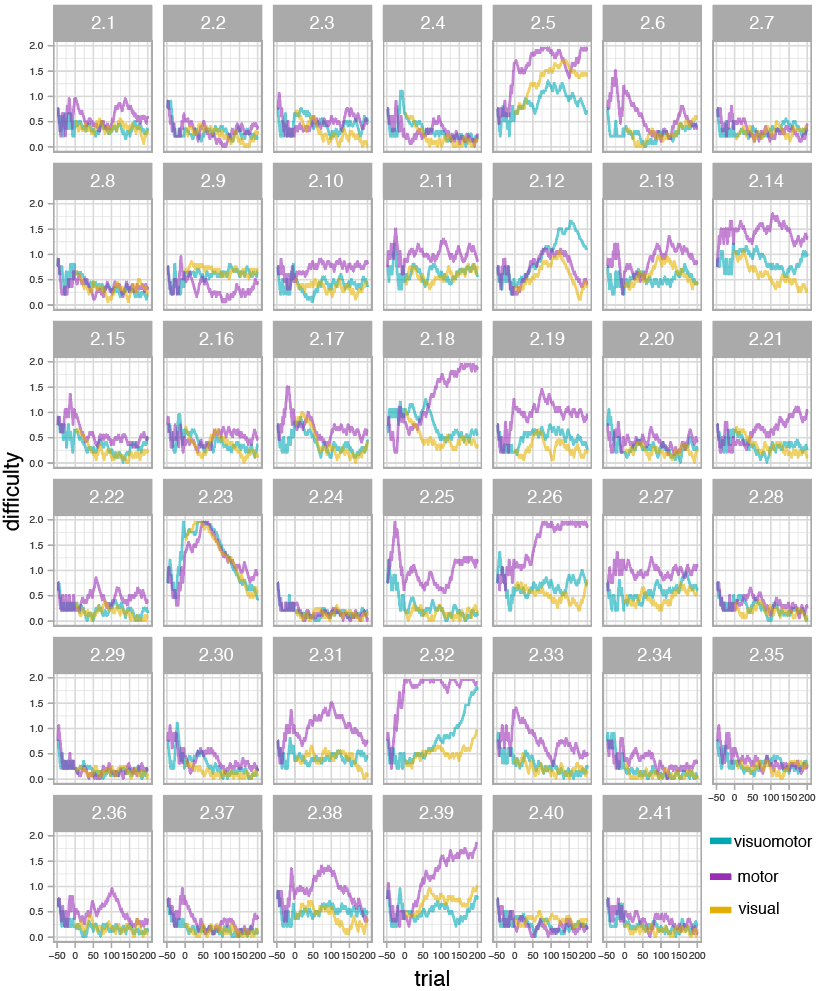

**Figure S2. *Staircasing procedure in Experiment 2, Skittles Trajectories task.*** Difficulty is expressed as difference in velocity (m/s). Only visuomotor and motor conditions were used in the calibration period.

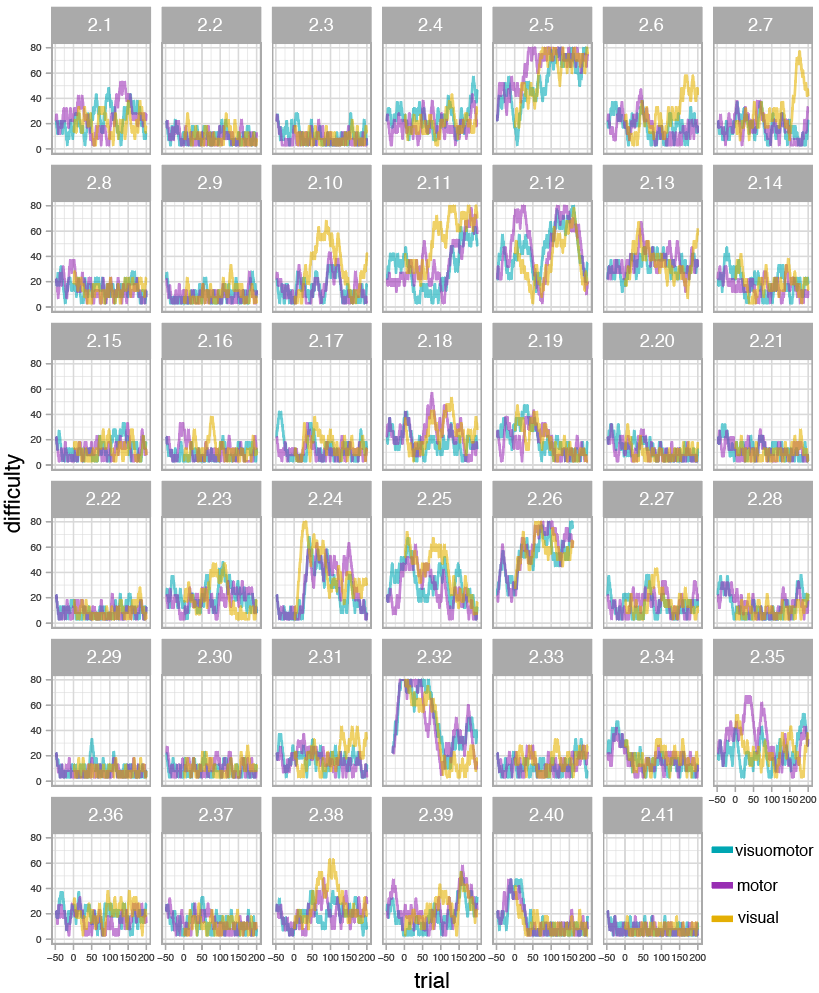

**Figure S3.** *Staircasing procedure in Experiment 2, Skittles Angles task.* Difficulty is expressed as a difference in degrees of angles. Only visuomotor and motor conditions were used in the training period.
